# A Willow Sex Chromosome Reveals Convergent Evolution of Complex Palindromic Repeats

**DOI:** 10.1101/710046

**Authors:** Ran Zhou, David Macaya-Sanz, Craig H. Carlson, Jeremy Schmutz, Jerry W. Jenkins, David Kudrna, Aditi Sharma, Laura Sandor, Shengqiang Shu, Kerrie Barry, Gerald A. Tuskan, Tao Ma, Jianquan Liu, Matthew Olson, Lawrence B. Smart, Stephen P. DiFazio

**Affiliations:** Department of Biology, West Virginia University, Morgantown, WV 26506-6057; Horticulture Section, School of Integrative Plant Science, Cornell University, New York State Agricultural Experiment Station, Geneva, NY 14456; HudsonAlpha Institute of Biotechnology, Huntsville, Alabama, USA; Department of Energy Joint Genome Institute, Walnut Creek, California, USA; Arizona Genomics Institute, School of Plant Sciences, University of Arizona, Tucson, AZ, USA; Biosciences Division, Oak Ridge National Laboratory, Oak Ridge, TN 37831, USA; DOE-Center for Bioenergy Innovation (CBI), Oak Ridge National Laboratory, Oak Ridge, TN 37831, USA; Key Laboratory of Bio-Resource and Eco-Environment of Ministry of Education, College of Life Sciences, Sichuan University, Chengdu 610065, China; State Key Laboratory of Grassland Agro-Ecosystem, Institute of Innovation Ecology & College of Life Sciences, Lanzhou University, Lanzhou 730000, China; Department of Biological Sciences, Texas Tech University, Box 43131, Lubbock, TX 79409- 3131, USA

**Keywords:** Sex, gene conversion, W chromosome, palindrome, genome, *Salix*

## Abstract

**Background:** Sex chromosomes in a wide variety of species share common characteristics, including the presence of suppressed recombination surrounding sex determination loci. They have arisen independently in numerous lineages, providing a conclusive example of convergent evolution. Mammalian sex chromosomes contain multiple palindromic repeats across the non-recombining region that facilitate sequence conservation through gene conversion, and contain genes that are crucial for sexual reproduction. Plant sex chromosomes are less well understood, and in particular it is not clear how coding sequence conservation is maintained in the absence of homologous recombination.

**Results:** Here we present the first evidence of large palindromic structures in a plant sex chromosome, based on a highly contiguous assembly of the W chromosome of the dioecious shrub *Salix purpurea*. Two consecutive palindromes span over a region of 200 kb, with conspicuous 20 kb stretches of highly conserved sequences among the four arms. The closely-related species *S. suchowensis* also has two copies of a portion of the palindrome arm and provides strong evidence for gene conversion. Four genes in the palindrome are homologous to genes in the SDR of the closely-related genus *Populus*, which is located on a different chromosome. These genes show distinct, floral-biased expression patterns compared to paralogous copies on autosomes.

**Conclusion:** The presence of palindromic structures in sex chromosomes of mammals and plants highlights the intrinsic importance of these features in adaptive evolution in the absence of recombination. Convergent evolution is driving both the independent establishment of sex chromosomes as well as their fine-scale sequence structure.

## Introduction

Sex chromosomes are fascinating not only because they contain the master regulatory genes that confer sex-specific traits, but also because of their unique evolutionary history [1]. In theory, the heterogametic (sex-specific) sex chromosome evolved from an autosome. There are two important features in sex determination regions (SDRs): suppressed recombination and the presence of sequences that only occur in one sex [1]. Furthermore, many sex chromosomes have lost most of their original genes over evolutionary time, and accumulated repetitive sequences such as transposable elements and tandem gene duplications [2, 3]. Consequently, ancient sex chromosomes are difficult to sequence because they are often highly heterochromatic and have a large amount of repetitive and ampliconic DNA [1, 4].

A striking characteristic of animal Y chromosomes is the presence of large palindromes in Y-ampliconic regions consisting of large inverted repeats with highly identical sequences that are undergoing gene conversion [5]. These palindromes are enriched for genes that are expressed in the testes and are important for male sexual development in mammals [4,6–8]. Palindromes have also been found on the W chromosomes of New World sparrows and blackbirds, suggesting that this may be a widespread feature of sex chromosomes [9]. However, such structures have not yet been described in plants.

Plant sex chromosomes have been identified in approximately 40 species but fewer than half of them are heteromorphic (readily discernable by cytology) [10, 11]. In plants, male-specific Ys (MSYs) have been studied in papaya and persimmon. Both of these contain a female suppressor on the Y chromosome [12–14]. Recently, a female suppressing gene in asparagus has been identified on the Y chromosome using long-read sequencing technology with optical mapping [15]. Another study on octoploid strawberry found repeated translocations of a female-specific gene cassette [16]. RNA-seq was applied to study the switches in heterogamety in two *Silene* species as well [17]. These dynamic sexual systems provide an excellent opportunity to study the drivers of sex chromosome evolution.

Sex determination is similarly diverse within the Salicaceae family. SDRs have been consistently found on chromosome 15 with female heterogamety in multiple *Salix* species [18–20]. This is quite different from the closely-related genus *Populus* where sex determining regions consistently occur on chromosome 19, with most species showing male heterogamety [21, 22]. Previously, we reported that the SDR occupies a large portion of the W chromosome in *S. purpurea*, because a large region on Chr15 shows lack of recombination and a large number of female-biased alleles [18, 23]. This is different from the small SDR observed in *P. trichocarpa* and *P. balsamifera* [22, 24]. However, due to the structural complexity of the SDRs, none of these studies have thus far included an in-depth analysis of the sequence composition and structure of the SDRs, and it is unclear whether there is a common underlying mechanism of sex determination. Here we present a much more complete assembly of the *S. purpurea* W chromosome and report for the first time in plants a palindromic repeat structure that is similar to the one found on mammalian Y chromosomes. We also demonstrate that shared homologous genes occur in the *Salix* and *Populus* SDRs, suggesting a potentially shared underlying mechanism of sex determination in this family.

## Results

### Genome assembly

We present here highly contiguous genome assemblies of a female and a male *S. purpurea*. The female assembly (94006 v4) consists of 452 contigs with an N50 of 5.1 Mb, covering a cumulative total of 317.1 Mb. Similarly, the male assembly (Fish Creek v3), has 351 contigs and an N50 of 5.6 Mb, covering 312.9 Mb (Additional File 1: Table S1). Both assemblies are partially phased in genomic regions where the two haplotypes are divergent. Alternative haplotypes are represented by 421 contigs totaling 72.4 Mb in the female assembly, and 497 contigs totaling 149 Mb for the male. Using a genetic map from a large family derived from progeny of the sequenced male genotype, we created assemblies representing the 19 chromosomes, containing 108 contigs totaling 288.3 Mb for the female, and 96 contigs totaling 288.5 Mb for the male. These represent over 90% of the assembled sequence in both cases, though 344 and 255 contigs remained unplaced by the genetic map for the female and male, respectively (Additional File 1:Table S2).

Because we expected the female-specific haplotype to be differentiated from the shared haplotype in the SDR, we anticipated that much of this region would be assembled as separate contigs, corresponding to W and Z haplotypes in the SDR. These can be readily differentiated by examining the relative depth of coverage when aligning male versus female short read sequences against these references. After identifying the location of the SDR based on the presence of sex-biased markers [18], the initial Chromosome 15 assembly appeared to consist of a mix of Z and W scaffolds in the SDR (Additional File 2: Figure S1a). We therefore sought to create a new assembly with Z and W haplotypes assembled to separate chromosomes. To do this we first identified the female-specific contigs (putative W contigs) using sex association and sex-specific depth as criteria. This resulted in identifying 23 contigs that were putatively comprised primarily of female-specific sequence (Additional File 1: Table S3). One scaffold was excluded because it mostly consisted of an alternative haplotype of a longer contig in the PAR of Chr15W.

Many of these contigs lacked markers from our intercross map, particularly for those that came from female-specific W (FSW) regions that were absent on the Z chromosome. We therefore created new genetic maps that had a mix of SNP and indel markers that would be more suited to capturing these hemizygous portions of the genome. The new genetic maps converged to 19 major linkage groups representing the 19 chromosomes. The male backcross map contained 8,715 markers, while the female backcross map contained 8,560 markers (Additional File 1: Table S4). We used these to assemble a Z and a W version of Chr15 (Additional File 1: Table S5). Thus, the current assembly (release ver5) contains 20 chromosomes, including Chr15Z and Chr15W. A total of 6.56 Mb (95.7%) of the W-specific contig sequence, contained in 17 contigs, was assembled to Chr15W using these maps. Four putative W scaffolds totaling 297 kb in length lacked mapped markers and could not be placed unambiguously.

### Location of the SDR

We repeated sex association analysis with our new assembly with Chr15Z removed. Among 54,959 tested SNPs, all 105 significantly sex-associated SNPs were present only on Chr15W (Fig. 1a; Additional File 2: Figure S2a-c), and markers from PARs and other scaffolds in the main genome did not show any sex association (Additional File 2: Figure S2a). The eight top-ranking sex-associated markers were distributed from 7.66 Mb to 8.66 Mb. Sex-associated markers were primarily heterozygous in females and homozygous in males, confirming our previously-reported observation of ZW sex determination in *S. purpurea* [18].

**Figure 1.**
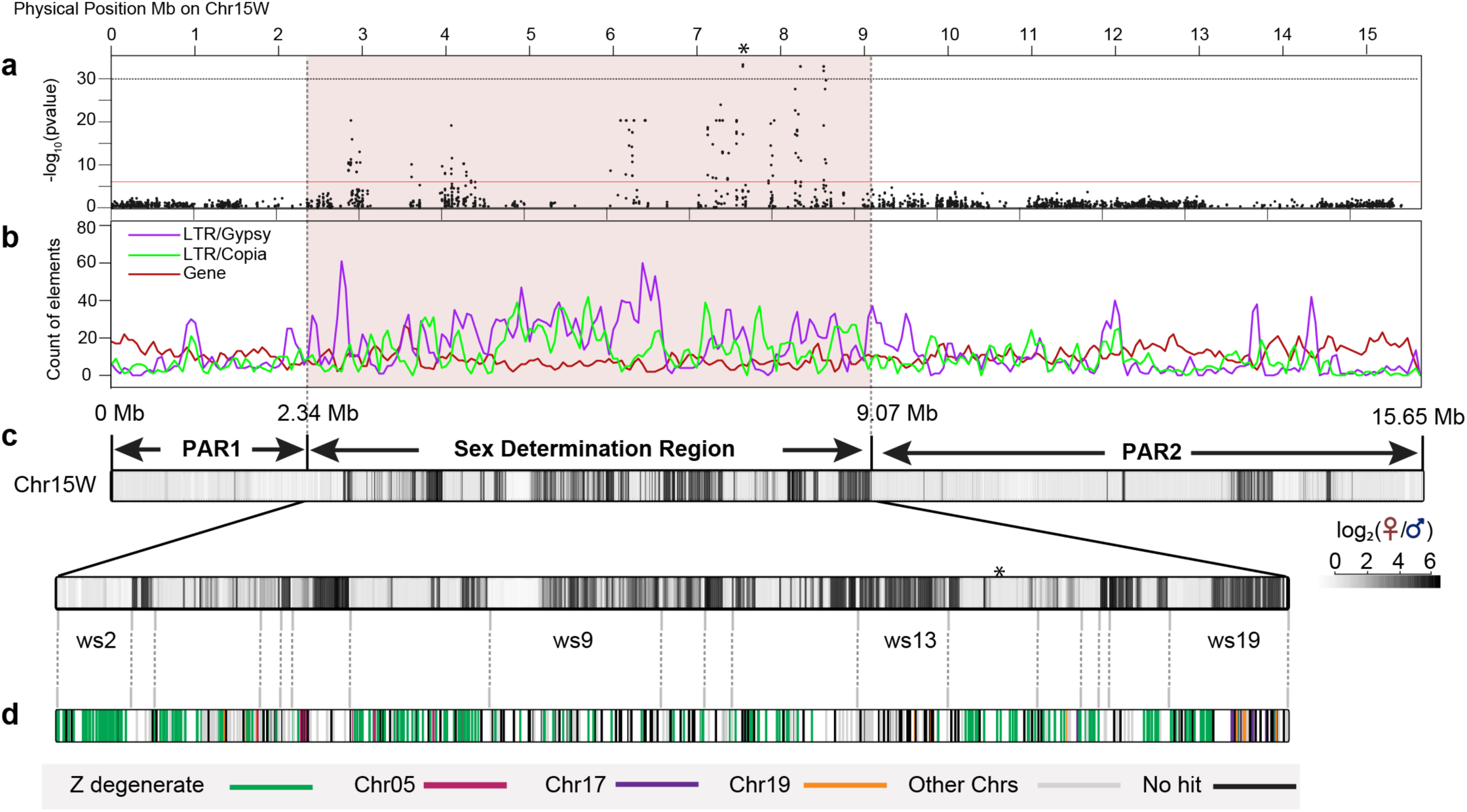
Genomic content of Chr15W and composition of the sex determination region (SDR). **a.** A Manhattan plot of Chr15W, based on GWAS using SNPs derived from aligning to a reference genome lacking Chr15Z. The Y axis is the negative logarithm of p values, and the red line indicates the Bonferroni cut off. **b**. Count of LTR elements including Gypsy and Copia, as well as genes in 100 kb windows with a 50 kb step size. **c**. Distribution of female-biased sequence on Chr15W, along with a more detailed view of the SDR below. Each colored block shows the log_2_ of the ratio of female and male depth in 10 kb windows. Vertical gray lines below the figure show the boundaries of the contigs in the SDR. **d**. Each tick represents a gene in the SDR. Colors indicate putative origins of the genes based on blastp versus the rest of the genome.

### Composition of the sex chromosomes 15W and 15Z

Chr15W is 15.7 Mb in length, composed of 22 contigs placed with the new genetic map. For comparison, Chr15Z is only 13.3 Mb and is comprised of 16 contigs (Additional File 1: Table S5; Figure 1). There are two pseudoautosomal regions (PARs) at each end of Chr15W that are indistinguishable from the corresponding regions on Chr15Z. PAR1 is 2.7 Mb long and is composed of two contigs, and PAR2 is 6.5 Mb and is comprised of three contigs (Fig. 1). These regions are unphased and are therefore identical in the two assemblies.

The sex-determining region (SDR) spans from 2.7 Mb to 9.1 Mb on Chr15W and occupies nearly 40% of the chromosome (hereafter referred to as the W-SDR). This region undergoes minimal recombination in the mapping population (Additional File 2: Figure S3). Reexamining sex-specific depth of the W-SDR, it is clear that this region of the genome is mostly phased to separate the male and female haplotypes (Additional File 2: Figure S1b). One contig (ws9) in the W-SDR is clearly chimeric based on the distribution of female-specific sequence (Fig. 1c), and the depth profile (Additional File 2: Figure S1b). However, this contig is only 300 kb in length, so it doesn’t change the overall conclusion that the FSW is largely phased. The region corresponding to the W-SDR on Chr15Z is about 4 Mb in length, and only occupies 28.2% of the chromosome (hereafter referred to as the Z-SDR). The size discrepancy is primarily due to the FSW regions. Based on the depth ratios, the Z-degenerate regions are about 3.5 Mb and W insertions are about 3.1 Mb in the W-SDR.

The W-SDR has lower gene density and higher repeat density than other portions of the genome (Table 1). More specifically, both the W-SDR and the Z-SDR show lower gene density on average than the PARs or other autosomes. Similarly, both the W-SDR and Z-SDR show higher accumulation of Gypsy retrotransposons. Interestingly, Copia-LTRs occur at higher density in the W-SDR region compared to the Z-SDR (15 per 100 kb vs 9 per 100 kb), (Kruskall-Wallis test, P<2.2e-16) (Table 1), suggesting that these inserted following cessation of recombination between these haplotypes.

**Table 1.** Density of genes and LTR retrotransposons in different areas of the genome. Numbers in parentheses are standard deviations of frequency per 100 kb window.

| Count per 100 kb | SDR | Z-SDR | PAR | Autosomes* |
| --- | --- | --- | --- | --- |
| Genes | 7.7(3.9) | 7.3(3.4) | 12.0(4.2) | 11.4(4.2) |
| Gypsy-LTR | 20.7(12.9) | 21.8(13.4) | 8.3(9.1) | 10.0(11.0) |
| Copia-LTR | 15.0(10.0) | 9.0(6.7) | 6.9(5.4) | 7.9(7.0) |

\* All 18 chromosomes are included.

### Gene content of the W chromosome

There are 269 genes in PAR1, 778 genes in PAR2, and 488 genes in the W-SDR. In contrast, the Z-SDR only contains 317 genes (Fig. 2; Additional File 1: Table S6-7). An additional 29 genes are present on scaffold_844, which is likely derived from the Z haplotype, but which lacked genetic markers to properly place it. There were 146 single copy mutual best hits between the W-SDR and Z-SDR, referred to hereafter as Z degenerate genes (Fig. 2). The SDR also contains 47 genes in tandem duplications, while the corresponding tandem repeats in the Z-SDR contain 78 genes. Additionally, the Chr15W SDR contains 65 genes that have mutual best hits on other autosomes, and 33 of these are tandemly duplicated in the SDR. In contrast, the Z-SDR region contains only 14 such genes, only four of which are tandemly duplicated. These putatively translocated genes comprise 13% of the W-SDR and only 4% of the Z-SDR, and account for nearly half of the discrepancy in gene content between the haplotypes. An additional 52 genes in the W-SDR had a top hit to other genes in the genome, but the best hit was not mutual, so these are lower confidence candidates for translocations or Z-degenerate genes. The Z-SDR contained 20 such genes. The remaining genes had no significant hits to other genes in the genome, presumably due to loss by deletion, or gaps in the sequence or annotation (125 in the W-SDR and 58 in the Z-SDR).

**Figure 2.**
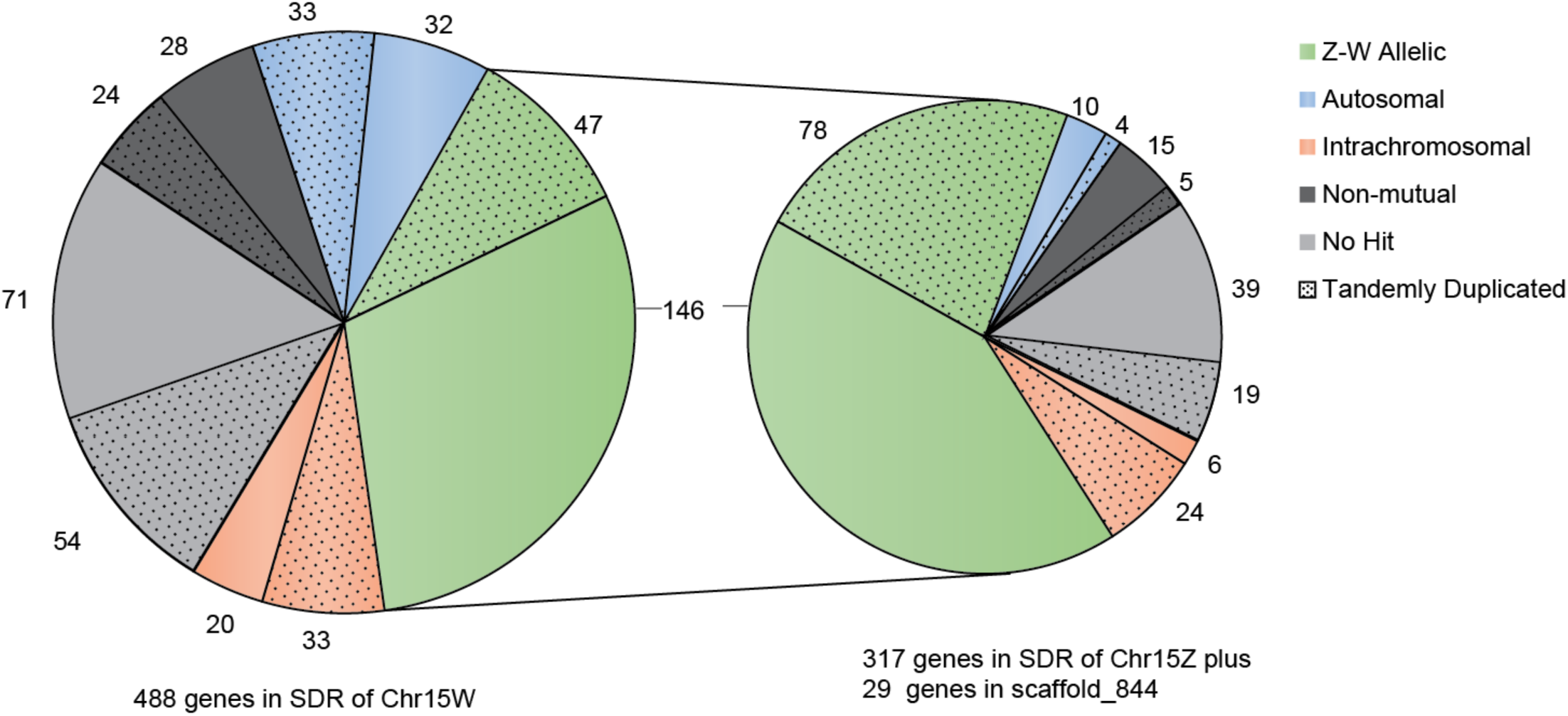
Annotated genes in Chr15W and Chr15Z. Genes are grouped according to the best non-self-hit in the annotated genome. Twenty-nine genes from an unmapped Z, scaffold_844 are also included. Stippled areas indicate genes of groups identified as tandem duplicates.

### Z-degenerate Genes and Strata

We used syntenic gene pairs identified through MCScanX between Chr15W and Chr15Z to test if there are strata with different levels of synonymous substitutions (Ks), which would indicate different phases of cessation of recombination [25]. There was little evidence to support the presence of strata based on 151 pairs of Z-degenerate genes (Fig. 3 and Additional File 1: Table S8). The average of Ks was 0.028. For comparison, the Ks between syntenic genes on Chr01 for *S. purpurea* and *S. suchowensis* was 0.027, and the Ks between *S. purpurea* and *P. trichocarpa* was 0.156 (Fig. 3).

**Figure 3.**
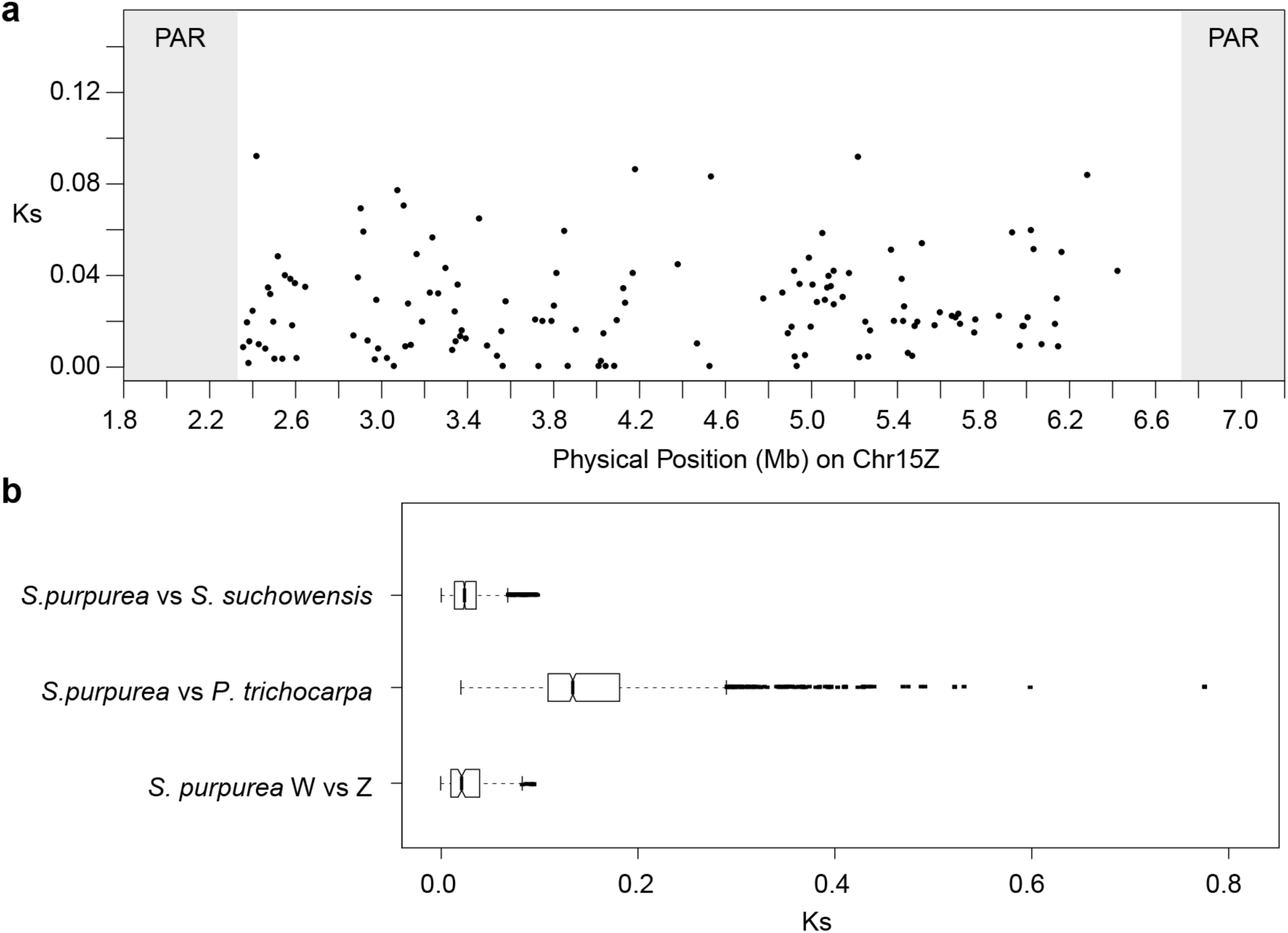
Synonymous substitution rates (Ks) for genes in the SDR. a. Comparison of syntenic genes in the W-SDR and Z-SDR. **b**. Boxplot showing distributions of interspecific synonymous substitutions for 1,203 syntenic genes on Chr01 for the closely-related species *S. purpurea* and *S. suchowensis* and for 1,377 genes on Chr01 in *S. purpurea* and *Populus trichocarpa*, compared to the distribution of substitutions between syntenic genes in the *S. purpurea* SDR.

### Translocations to the W-SDR and palindromic repeats

The recently translocated genes are of particular interest because of the insights they can provide into the evolutionary history of the sex chromosome. Among 65 genes putatively translocated from autosomes to the W-SDR, 10 have best hits on Chr19 (manually annotated genes excluded) (Additional File 1: Table S9). Contig ws19 is particularly enriched for translocated genes, and merits a closer examination (Fig. 1). Contig ws19 contains 11 translocated genes, including four genes from Chr19 and four genes from Chr17 (Fig. 1). Many of these translocated genes occur in two to four copies on ws19 in striking inverted repeat configurations that are similar to the palindromic repeats that occur on mammalian Y chromosomes (Fig. 4).

**Figure 4.**
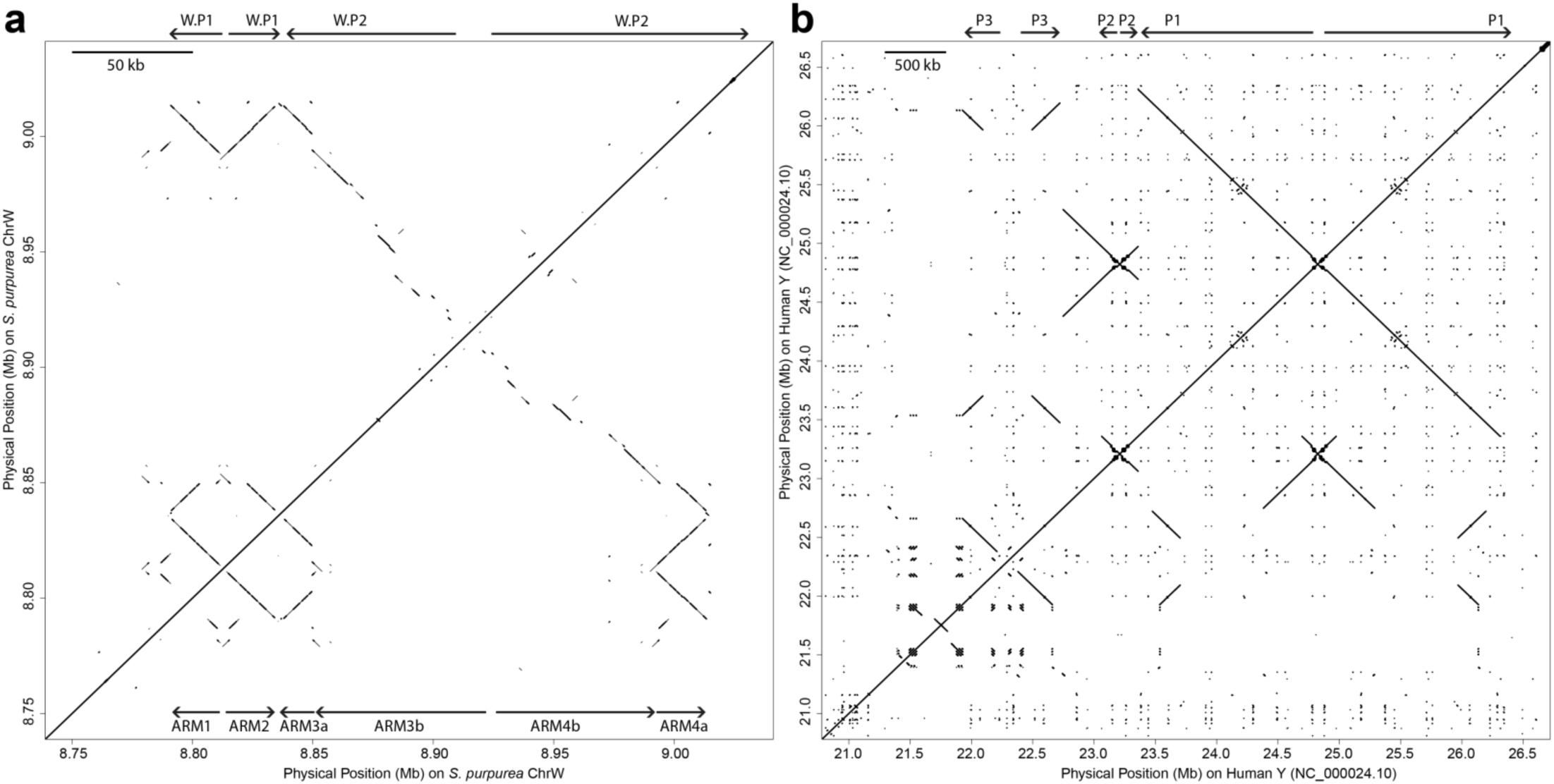
Palindromic repeats in the *S. purpurea* W chromsosme (a) and the *H. sapiens* Y chromosome (b). The dot plots were produced using LASTZ with identical settings. Note the different scales, indicated by the bar at the top right of each figure. *H. sapiens* palindromes are labeled following Skaletsky et al. [4].

In *S. purpurea*, this region is female-specific and is composed of two palindromes. Palindrome W.P1 spans about 42.7 Kb with a 2.6 kb spacer in the center, and Palindrome W.P.2 is immediately adjacent and spans over 165 kb (Table 2; Fig. 4a). A 20 kb sequence occurs in inverted orientation and shows high sequence identity across the four arms of both palindromes (Table 2; Fig. 5a). In palindrome W.P1 these are referred to as arm1 and arm2, and in Palindrome W.P2 these are referred to as arm3a and arm4a (Table 2; Fig. 4a). Sequence identity among these four arms is greater than 99% on average. The regions of high identity are disrupted by a ∼500 bp insertion in the center of arm4. Furthermore, arm3 has a 6.9 kb deletion at 11.7 kb, followed by a stretch of 1.6 kb that can be aligned to the other arms in the same orientation (Fig. 5a). Additionally, there is a 12 kb stretch upstream of arm1 that shows high identity to portions of arms 1 and 2. We call this the pre-arm for convenience (Table 2).

**Figure 5.**
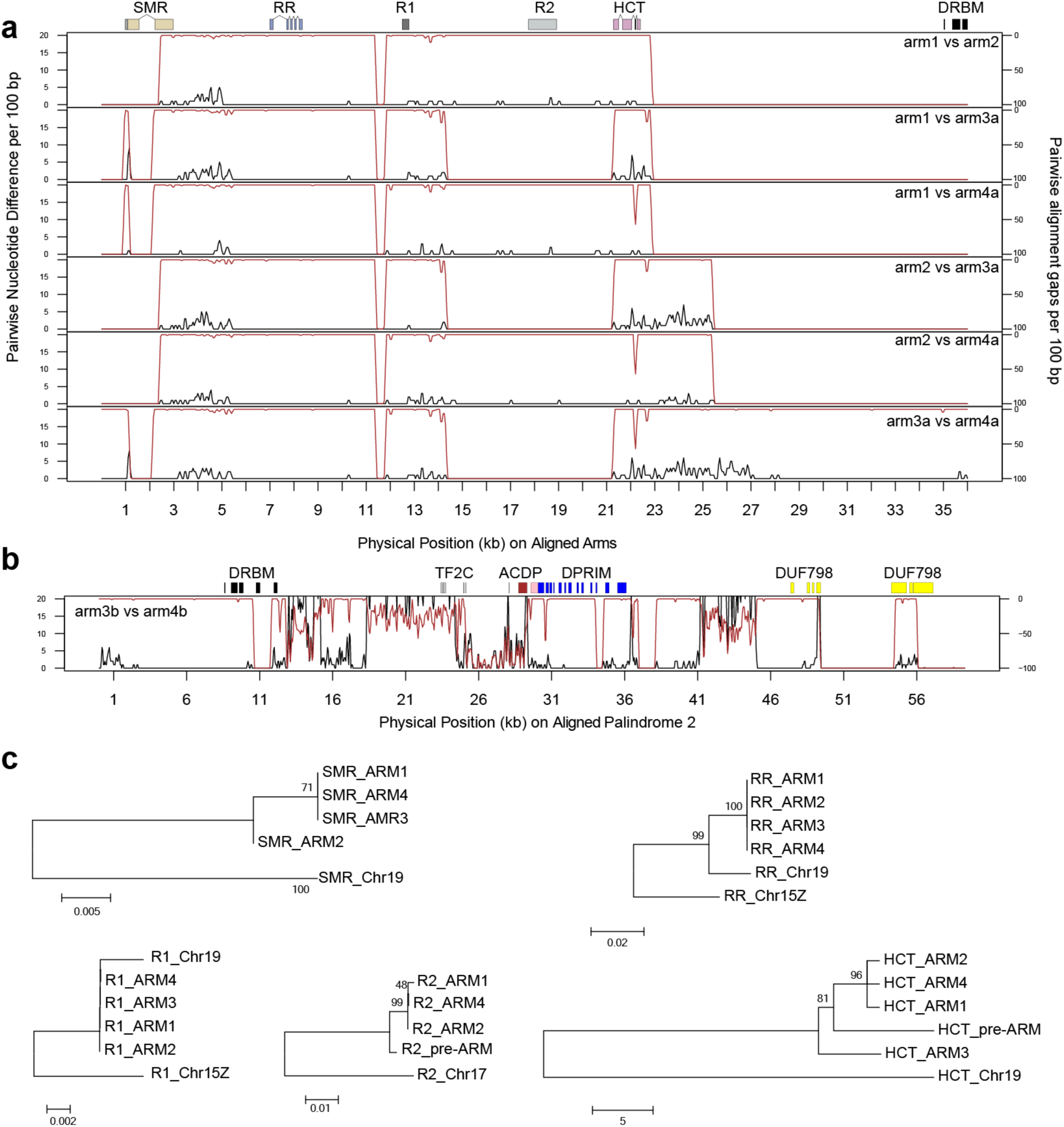
Sequence comparisons for the two palindromes. a. Comparison of the four arms that are shared among the two palindromes. The black line represents the number of nucleotide differences in 100 bp windows, while the red line indicates gaps in the alignment on an inverted scale. b. Comparison of the portions of palindrome 2 that are not shared with palindrome 1. c. Phylogenetic trees of five multi-copy genes in the palindromic region.

**Table 2.** Coordinates of palindromes in the female SDR.

|  | name | Start (bp) | End (bp) | Size (bp) | Gene families |
| --- | --- | --- | --- | --- | --- |
| Palindrome W.P1 | pre-arm | 8,778,973 | 8,791,042 | 12,070 | *R2,HCT* |
|  | arm1 | 8,790,932 | 8,811,002 | 20,071 | *SMR,RR,R1,R2,HCT* |
|  | Spacer1 | 8,811,003 | 8,814,588 | 3,586 |  |
|  | arm2 | 8,814,589 | 8,834,138 | 19,550 | *SMR,RR,R1,R2,HCT* |
| Palindrome W.P2 | arm3a | 8,836,813 | 8,850,772 | 13,960 | *SMR,RR,R1,HCT* |
|  | arm3b | 8,850,773 | 8,920,527 | 69,755 | *DRBM,TF2C,DPRIM,DUF789* |
|  | Spacer2 | Unidentified |  |  |  |
|  | arm4b | 8,920,528 | 8,993,098 | 72,571 | *DRBM,ACDP,DPRIM,DUF789* |
|  | arm4a | 8,993,099 | 9,013,390 | 20,292 | *SMR,RR,R1,R2,HCT* |

Palindrome W.P2 contains an additional inverted repeat that is missing from W.P1. We refer to this as arm3b and arm4b (Table 2; Fig. 4a). Sequence conservation is somewhat lower between these two arms compared to the other four, ranging from 96% to 99% over most of their length. Furthermore, the regions of high identity are disrupted by numerous insertions and deletions (Fig. 5b).

### Gene content of the palindromes

There are five genes duplicated across arms 1, 2, 3a and 4a of both palindromes. These are the Small Muts-Related protein (SMR), a Type-A cytokinin response regulator (RR), two genes that contain an NB-ARC domain (R1 and R2), and a Hydroxycinnamoyl-CoA shikimate/ hydroxycinnamoyl transferase (HCT) (Table 3). All of these genes except R2 have clear paralogous copies on Chr19. There is very little sequence divergence among most of these paralogs in the palindromes (Fig. 5).

**Table 3.** Genes present in Palindromes 1 and 2.

|  | Gene Symbol | Number of Copies | GeneID | Chromosome of the non-W best hit | Best Hit in *A. thaliana* | Arabidopsis name or description (function) | Best Hit in *P. trichocarpa* v3 | Identity of *P. trichocarpa* Best Hit |
| --- | --- | --- | --- | --- | --- | --- | --- | --- |
| Palindromes W.P1 and W.P2 | *SMR* | 4<sup>[a]</sup> | Manually annotated | Chr19 | AT5G23520 | *SMR* (Small MutS Related domain-containing protein) | Potri.T013000 | 90.70 |
|  | *RR* | 4 | Sapur.15WG073500<br>Sapur.15WG073900<br>Sapur.15WG074000<br>Sapur.15WG075200 | Chr19 | AT3G56380 | *ARR17* (type A cytokinin response regulator) | Potri.019G133600 | 92.81 |
|  | *R1* | 4<sup>[a]</sup> | Sapur.15WG073800<br>Sapur.15WG074100 | Chr15Z | AT4G27220 | NB-ARC domain-containing disease resistance protein | Potri.T012900 | 81.00 |
|  | *R2* | 3(1)<sup>[a]</sup> | Manually annotated | Chr17 | AT4G27220 | NB-ARC domain-containing disease resistance protein | Potri.T013300 | 61.23 |
|  | *HCT* | 4(1) | Sapur.15WG073400<br>Sapur.15WG073600<br>Sapur.15WG073700<br>Sapur.15WG074200<br>Sapur.15WG075100 | Chr19<sup>[b]</sup> | AT5G48930 | *HCT* (hydroxycinnamoyl-coa shikimate/quinic acid hydroxycinnamoyl transferase) | Potri.018G104700 | 58.02 |
| Palindrome W.P2 only | *DRBM* | 2 | Sapur.15WG074300<br>Sapur.15WG075000 | Chr17 | AT1G09700 | *ATDRB1* (dsRNA binding protein) | Potri.017G126700 | 61.95 |
|  | *TF2C* | 1 | Sapur.15WG074400 | Chr08<sup>[c]</sup> | AT2G27040 | *AGO4* (*ARGONAUTE 4*, siRNA mediated gene silencing) | NA | NA |
|  | *ACDP* | 1 | Sapur.15WG074900 | Chr17 | AT5G52790<sup>[d]</sup> | CBS domain protein with DUF21 (transmembrane transporter) | Potri.017G147900 | 83.33 |
|  | *DPRIM* | 2 | Sapur.15WG074500<br>Sapur.15WG074800 | Chr17 | AT5G52800 | DNA primase | Potri.017G148000 | 92.52 |
|  | *DUF789* | 2 | Sapur.15WG074600<br>Sapur.15WG074700 | Chr17 | AT1G03610 | *DUF789* (protein of unknown function) | Potri.017G152600 | 86.03 |

<sup>[a]</sup> Manually annotated transcripts were included in the count. Numbers in the parenthesis are from a fragment in the upstream portion of W.P1 that is homologous to part of W.P1.
<sup>[b]</sup> This cluster of tandem duplications on Chr19 in *S. purpurea* is not present on Chr19 in *P. trichocarpa*.
<sup>[c]</sup> The palindrome gene contains only a truncated blast hit to Sapur.008G005800 on Chr08.
<sup>[d]</sup> This best hit with an expected value of $8 \times 10^{-3}$ due to a sequence length of 84 aa. Expected values of the remaining *A. thaliana* were less than $1 \times 10^{-10}$ .

The cytokinin response regulator is of particular interest because an ortholog of this gene has also been found to be associated with sex in *Populus* [22], and is therefore an excellent candidate as a master regulator of sex in the Salicaceae. The RR gene is highly conserved across all four palindrome arms on the W-SDR (Fig. 5a,c). Interestingly, we also found a pseudogene copy of the RR gene on the Z-SDR. This is the only one of the five genes that is present in some form on the W-SDR, the Z-SDR, Chr19, and also in the SDR of *Populus*. There is a 2.6 kb sequence inserted upstream of all RR copies in the palindrome, and not in the Z-SDR pseudogene or on Chr19 (Additional File 2: Figure S4). This suggests that the W-SDR palindrome formed after translocation from Chr19. Interestingly, the RR gene also occurs as inverted repeats in all three locations in the genome (W-SDR, Z-SDR, and Chr19). However, alignment of the W-SDR, Z-SDR and Chr19 versions demonstrates that the palindromes likely formed independently, because the palindromic regions are different (Additional File 2: Figure S4).

There are an additional five genes in the W.P2 palindrome. Three of these genes occur as inverted repeats: a DNA-directed primase/polymerase protein (*DRBM*), a DNA primase (*DPRIM*), and a protein containing Domain of Unknown Function 789 (*DUF789*). In addition, there is a homolog of ARGONAUTE 4 (*TF2C*) and a CBS domain protein (*ACDP*) in single copy. Four of these genes were apparently translocated from Chr17 (Table 3). This leads us to the hypothesis that after these genes were translocated to the W-SDR they underwent several rounds of structural rearrangements, including duplications, inversions, and deletions.

### Multiple LTR retrotransposons in the palindrome

To gain further insight into the composition and history of the W-SDR, we used LTRharvest and LTRdigest to annotate LTR retrotransposons in the palindromic region. We identified one LTR retrotransposon in the pre-Arm region and 12 LTR retrotransposons in palindrome W.P2 that have terminal repeats identified with coding regions (Fig.6a). These 13 retrotransposons are likely to be independent insertion events given that they have different long terminal repeats as well as different target site duplications and do not occur in the same position in the opposite arm of the palindrome (Additional file 1: Table S10). Given that there are varying numbers of substitutions within the LTRs of the same retrotransposon, it appears that these insertions have occurred repeatedly after establishment of the palindromes. Using a previous estimation of the mutation rate in *P. tremula* (2.5×10^−9^ per year)[26], we estimate that the oldest insertion occurred at least 8.6 ± 2.9 s.d. MYA from a nonautonomous LTR retrotransposon, *Ltr-p2-a* (Fig. 6a and Additional file 1: Table S10). This is likely an underestimate, since the *Salix* substitution rate is substantially higher than that of *Populus* [27]. Since the oldest substitutions occurred in Palindrome W.P2, we infer that this element became established first (Fig. 6a). The LTRs of the nonautonomous elements *Ltr-p2-a* and *Ltr-p2-k* flank the *SMR* and *RR* genes (Fig. 6c,d; Additional file 2: Figure S5), which raises the intriguing possibility that these LTRs were involved in the translocation of these genes to this region. We also found two highly identical LTRs from the same family in W.P1 (*Ltr-p2-b3* on arm3 and the *Ltr-p2-b4* on arm4; Fig. 6a-c; Additional file 1: Table S10). There are truncated parts of this LTR in the pre-arm and the spacer between arm1 and arm2 as well (Fig. 6b, c). These copies might be a direct consequence of duplications and inversions that occurred during the formation of the palindrome instead of independent insertions.

**Figure 6.**
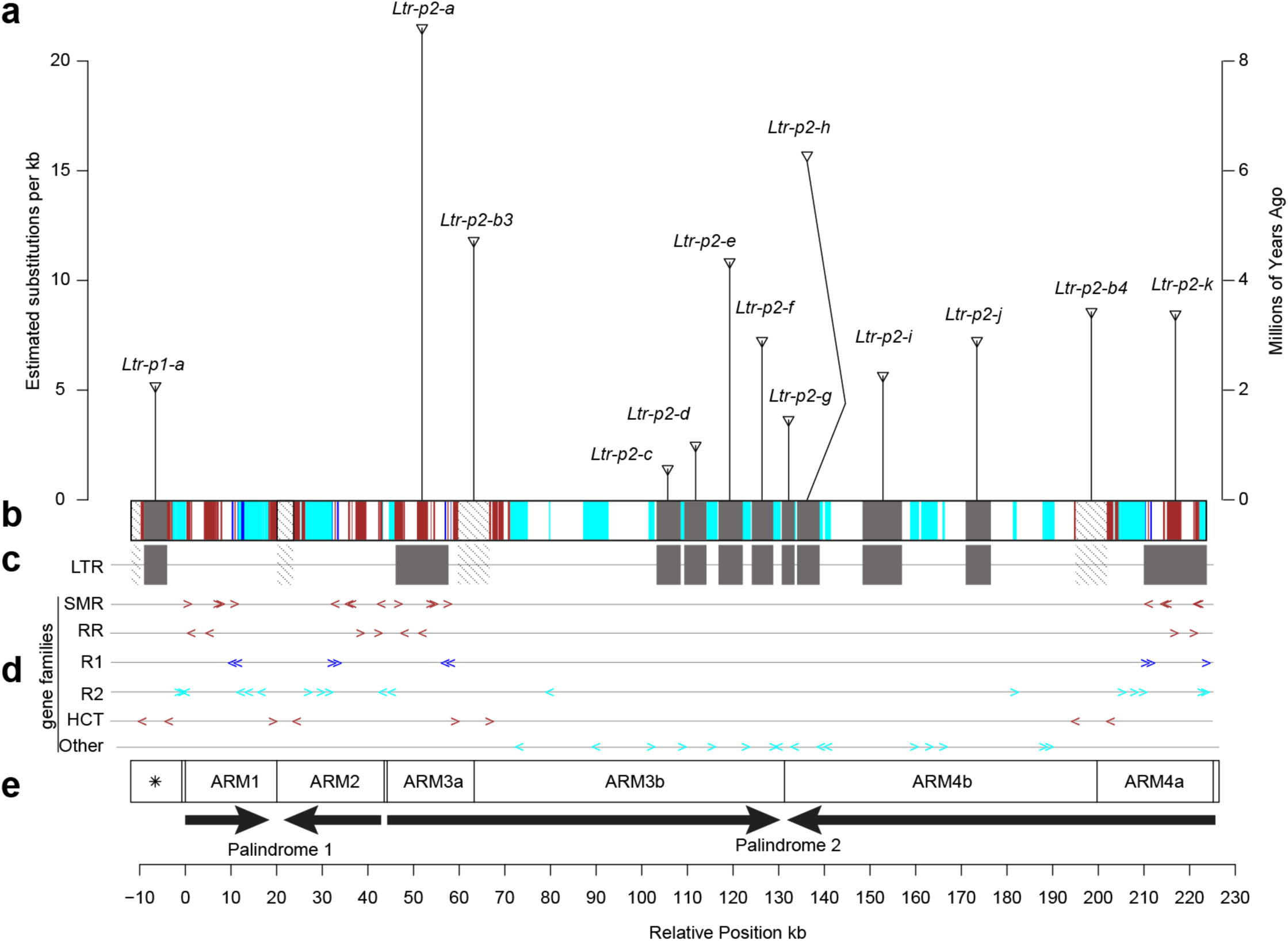
LTR retrotransposons, female specific genes, and palindromes. **a**. Each vertical line with a wedge on top represents each of the 13 TEs identified in the palindromic region by LTRharvest. The height of each line indicates the number of estimated nucleotide substitutions in the two LTRs (transposons a-h), and an approximation of the insertion time based on the mutation rate in *P. tremula* (Ingvarsson 2008). **b**. Colored boxes represent putative chromosomal origins of genes in the palindrome. Dark red, Chr19, cyan, Chr17. Blue boxes represent genes with paralogs on the Z chromosome. **c**. The positions of 13 LTRs (shaded boxes). Hatched boxes represent incomplete duplications derived from *Ltr-p2-b3/b4*. **d**. Exon positions and orientations, represented by colored arrows. **e**. Schematic representation of female-specific palindromes. The box with a star represents a homologous region derived from part of one of the arms (preARM). Directions of arrows indicate the relative orientations of the four arms.

### Evidence for gene conversion in the palindromes

We have shown that the palindromes are likely to be millions of years old, yet sequence identity of portions of the palindrome arms remains high (Fig. 5a). A likely explanation for this is gene conversion among the palindrome arms, as has been observed in the mammalian Y chromosome palindromes [5]. To test for this, we searched for regions that had interspecific polymorphisms relative to *S. suchowensis*, a closely-related species with ZW sex determination [20]. If regions with interspecific polymorphisms lack paralogous sequence variation (PSV) across the palindrome arms, then this would be excellent evidence of gene conversion. We detected a 3 kb region within the palindromes where there are no PSVs in *S. purpurea* and only one PSV in *S. suchowensis*, but substantial interspecific polymorphisms (Fig. 7). The depth of this region is 4N as expected for the four copies of the palindrome arms in *S. purpurea*. In *S. suchowensis*, the depth is between 2N and 3N, which indicates that there might be a palindrome structure as well, though it might be incomplete. We also applied the same methods with resequencing reads of two female and two male *S. viminalis* individuals (another *Salix* with ZW sex determination) [19], but the palindromic region was not well covered by reads of either sex. This may indicate that *S. viminalis* lacks the palindrome, though it is more distantly related to *S. purpurea* than is *S. suchowensis*, so this may simply be due to excessive sequence divergence in this region.

**Figure 7.**
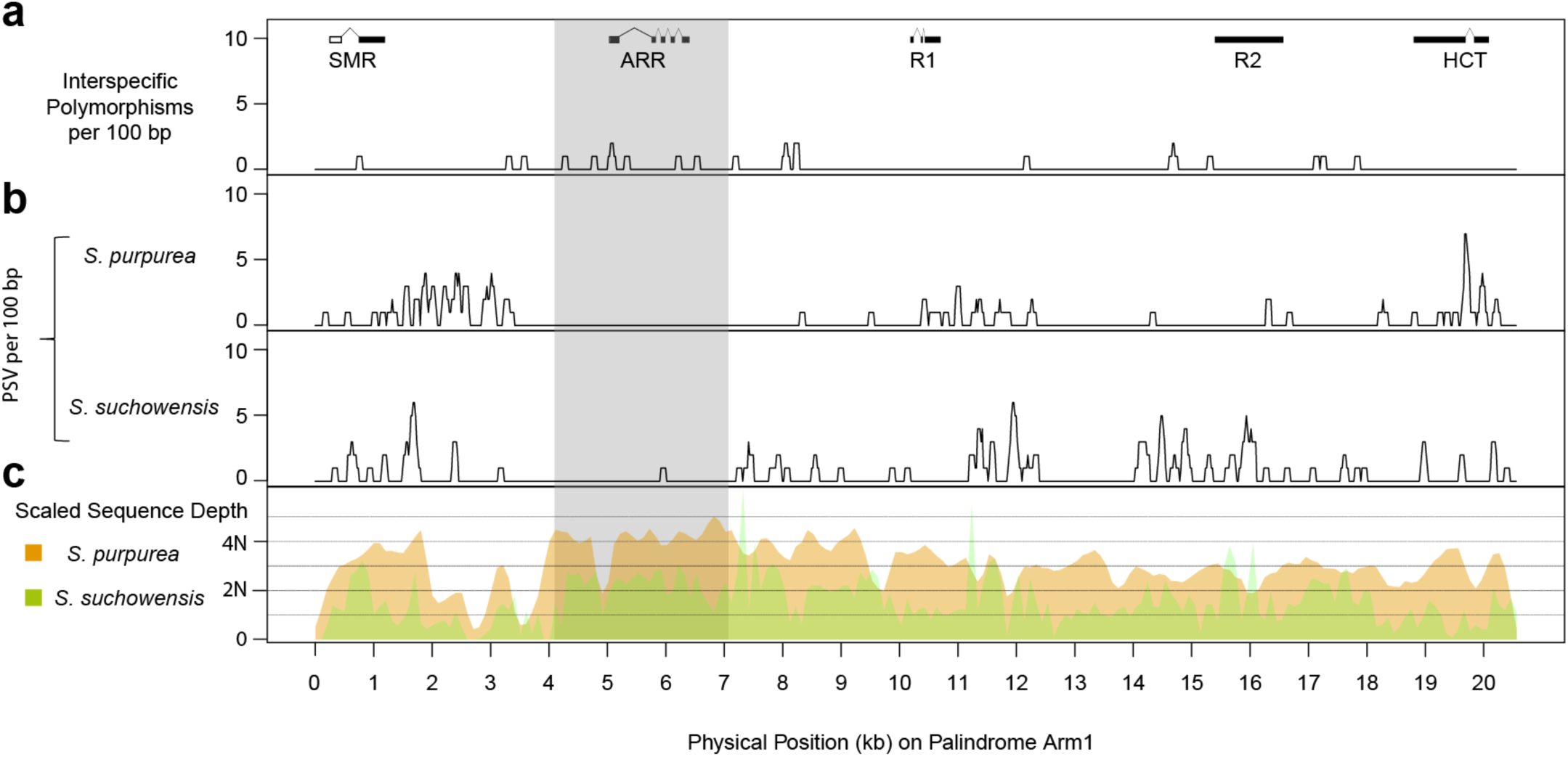
Sequence variation in the palindrome arms. **a**. Density of fixed differences between *S. purpurea* and *S. suchowensis* per 100 bp. **b**. Density of paralogous sequence variants (PSVs, differences among the 4 palindrome arms) in *S. purpurea* and *S. suchowensis*. **c**. Relative depth of Illumina sequence reads aligned to a reference sequence of one arm of the *S. purpurea* palindrome, where 2N represents the expected depth of read alignment across the whole genome. The grey shaded area represents a segment of the palindrome that is enriched for interspecific fixed variants, but depleted in PSVs, providing strong evidence for differential gene conversion in the two lineages.

### Expression patterns of genes in the palindromes

We examined expression profiles in multiple tissues of the two reference genomes to validate the predicted transcripts and to determine how the expression patterns of genes in the palindromes differ from their autosomal counterparts. As expected, most genes in the palindromes show female-limited expression while the autosomal copies are generally not sex-biased (Fig. 8a). The cytokinin response regulator (*RR*) (Sapur.15W073500) shows the highest expression in catkin tissue, followed by expression in shoot tips and stems. On the contrary, two autosomal copies on Chr19 show lower expression, limited to female catkins and male buds. The four copies of the *SMR* gene show low expression in female catkins and other tissues, but the autosomal copy on Chr19 (Sapur.019G001500) is expressed in all tissues (Fig. 8a). All five copies of the *HCT* gene from the palindromes showed low expression in female catkins and roots and higher expression in leaf tissues, shoot tips, and stems, all of which were female-biased. Two copies of the DNA Primase gene from palindrome W.P2 also show high expression in leaf tissues while the original copy on the autosome (Sapur.017G119600) was expressed across all sampled tissues. Similarly, analysis of transcriptomic data of catkins from 10 females and 10 males in the F_2_ family confirms that the genes in the palindromes are primarily expressed in female tissue, in contrast to their autosomal paralogs (Fig. 8b).

**Figure 8.**
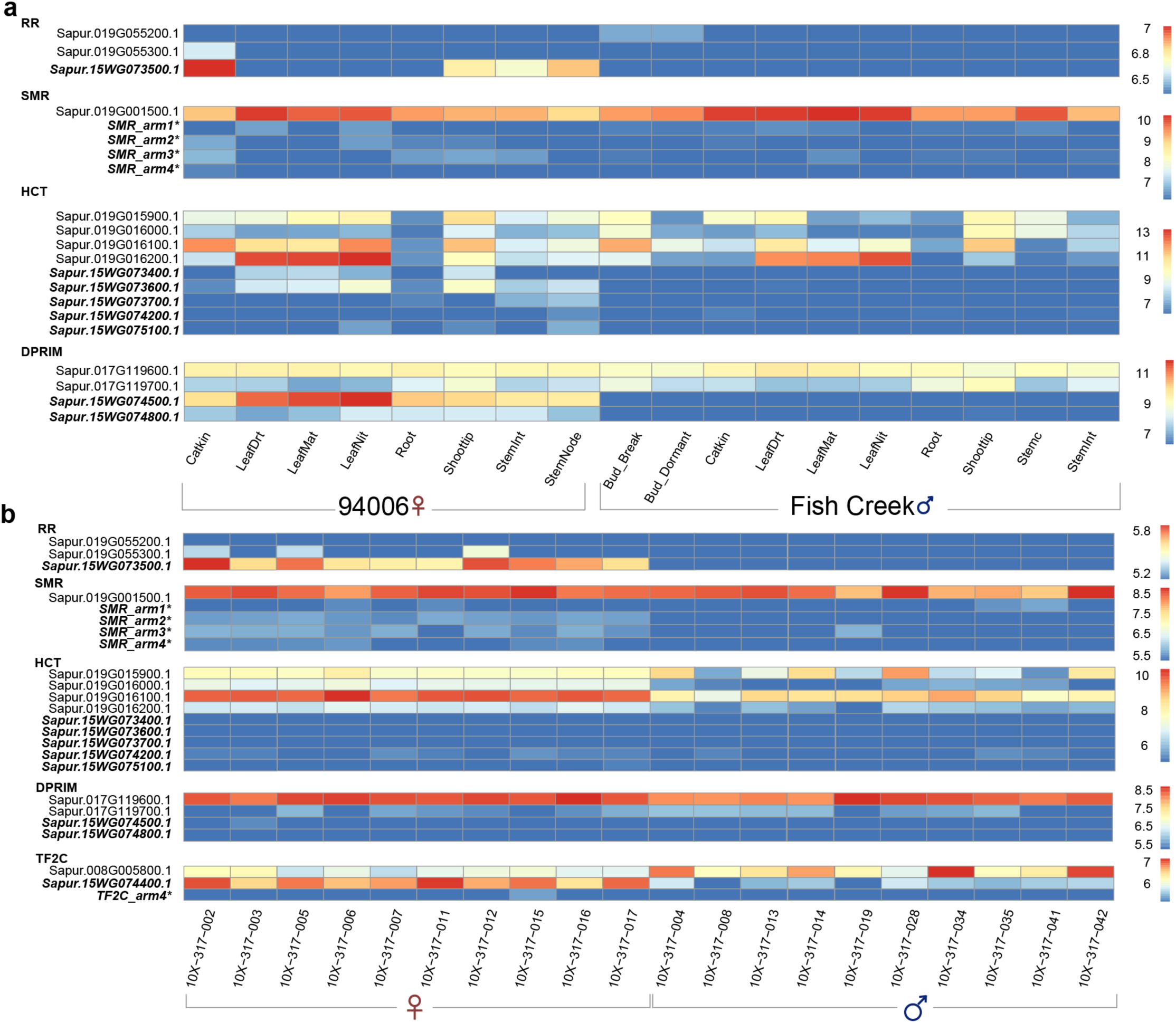
Expression profile of genes from the W palindromes and autosomal paralogs. **a**. Normalized read counts of genes in different tissues from clone 94006 (female) and Fish Creek (male). **b.** Normalized read counts of selected genes in catkins from 10 females and 10 males from an F_2_ family. Gene labels in bold font are from the palindromes. Asterisks indicates manually annotated genes.

## Discussion

### The W chromosome in *S. purpurea*

Using male and female specific depth from a controlled cross pedigree, we have been able to identify Z and W haplotypes from the SDR of a highly heterozygous species from a standard PacBio assembly. We also show how presence-absence markers generated from sequence depth in controlled cross progeny can be used to genetically map hemizygous portions of the SDR. In a similar study of a young Y chromosome in asparagus, BioNano optical maps for a YY individual were generated to improve genome contiguity, and sequence depth of coverage was also treated as a QTL to aid the assembly because of the presence of large indels in the sex chromosome [15]. Here, we showed that by combining long-read sequencing with GBS marker data from a large F_2_ family, we could efficiently identify the sex-specific regions of the sex chromosome. However, unlike strategies like single-haplotype iterative mapping and sequencing (SHIMS) that have been used in assemblies of mammalian Y chromosomes [4,6–8], our map-based strategy could not provide a definitive order for the female-specific contigs due to lack of recombination in the SDR.

Our W chromosome assembly has helped explain why sex association analyses sometimes detect multiple unlinked loci on different chromosomes [18,22,28]. GWAS with a nearly exhaustive collection of W contigs in the reference genome yields a single sex association peak (Additional File 2: Figure S2). We have shown that the SDR contains many translocations from autosomes. If those translocated segments are absent from the reference genome, W-specific reads from a female individual align to the original source of those sequences, resulting in false association peaks on the autosomes. This may also explain the multiple peaks detected in *P. trichocarpa* and *P. balsamifera*, both of which have XY sex determination systems. In this case, SNPs were identified based on alignments to a female reference genome, which lacks the SDR [22].

Autosomal translocations are likely to be important contributors to the evolutionary history of the sex chromosome in *S. purpurea*. The W chromosome is approximately 2.5 Mb larger than the Z chromosome due to expansion of the SDR, including a substantial expansion of gene content, driven in part by numerous translocations and subsequent expansion of autosomal genes. Autosomal translocations have also been demonstrated in other sex chromosomes. For example, ampliconic sequences on the human Y chromosome were acquired through transpositions from diverse sources, and then amplified [4]. These ampliconic sequences account for about 30% of the male-specific Y euchromatin [4]. These translocations and other structural rearrangements may also suppress recombination and cause linkage between autosomal and sex-linked genes [2]. In plants, the large sizes of young Y chromosomes have been attributed to the accumulation of retrotransposons (reviewed in [29]).

Initial analyses in *P. trichocarpa* suggested that the SDR is much younger than the whole genome duplication event that is shared by *Populus* and *Salix*, suggesting that the SDR became established well after these genera diverged [22]. The SDR of *S. purpurea* also appears to be established relatively recently. Furthermore, W and Z alleles in the nonrecombining SDR have similar Ks values compared to those for orthologous autosomal genes from *S. purpurea* and *S. suchowensis*. Given that the SDR is located in approximately the same portion of Chr15 in both species, and both have ZW systems [18, 20], it is reasonable to assume that the SDR became established in this lineage prior to divergence of these two species, but well after divergence from *Populus*.

Sex chromosomes commonly show evidence of “evolutionary strata” with markedly different Ks values that represent different epochs of expansion of the SDR [30]. Under one common model of sex chromosome evolution, these strata are the result of gradual SDR expansion as sexually antagonistic alleles become incorporated into the SDR [25, 31]. We detected little evidence for the existence of such strata in *S. purpurea*. This corroborates a previous analysis that failed to detect strata in *S. suchowensis* using an integrated segmentation and clustering method, which was successful in finding strata in multiple other species, including *P. trichocarpa* [32]. The failure of this method in *S. suchowensis* was hypothesized to be due to the small size of the SDR [32]. However, the identified SDR in *S. purpurea* is about 6-7 Mb, occupying more than one third of the W chromosome assembly. It appears instead that cessation of recombination has not been a gradual long-term process in the *S. purpurea* SDR. One explanation for the large size of this region is it partially overlaps with the centromere of Chr15, as we previously reported [18]. It’s possible that the repressed recombination in this region pre-dated the translocation of a relatively small SDR cassette, as has been observed in octoploid *Fragaria* [16]. This is consistent with the apparently small size of the region in *Populus* [22]. This is also consistent with the structure and composition of the palindromic repeats that we discovered in *S. purpurea*, as detailed below.

### Sex chromosome palindrome repeats

We have reported here the first observation of a large inverted repeat in a plant sex chromosome, similar to the palindromic structures observed in mammalian sex chromosomes. W.P1 and W.P2 of *S. purpurea* have a similar arrangement of arms as P1 and P3 in humans due to the presence of highly homologous regions between the two palindromes. Similar palindromes have been discovered on Y chromosomes of several mammals and avian W chromosomes (reviewed by [5]). Large mammalian palindromes developed as a series of accumulations of insertions from autosomes and maintained through arm-to-arm gene conversion. This intrachromosomal gene conversion provides an essential function in maintaining coding sequence integrity which otherwise would be compromised by the continuous accumulation of deleterious mutations in the absence of homologous recombination (i.e., Muller’s Ratchet) [5,33,34]. The fact that these structures have independently evolved in non-recombining regions of sex chromosomes is an intriguing case of convergent evolution of chromosome structure. It is interesting to note that the chloroplast genome, another non-recombining chromosome in plants, also contains a different large inverted repeat that undergoes gene conversion [35] and helps maintain structural integrity of the genome, suggesting that this phenomenon may be common in regions of the genome that lack recombination [36].

The *S. purpurea* palindromes are considerably smaller than mammalian palindromes, and have only accumulated two major autosomal translocations, possibly reflecting their young age. Another difference between the human palindrome and the one in *S. purpurea* is that the gene conversion seems to be quite efficient across all the eight palindromes in humans, but the observed regions under gene conversion in *S. purpurea* are much more limited. This is particularly obvious in W.P2, compared to human P1, which has high sequence identity over several Mb (Fig. 4).

LTR retrotransposons found in the palindrome are important at least for two reasons. The observation of nonautonomous LTR retrotransposons suggests that translocations mediated through transposable elements could provide a means of movement of genes (such as sex-determining gene(s)), similar to what has been observed in octoploid *Fragaria* [16]. Secondly, illegitimate intrachromosomal recombination events could be mediated by LTR retrotransposons that have accumulated in the SDR [37], resulting in complex rearrangements like the palindromic structures. Our observation of a truncated copy of a copia LTR located at the spacer between arm1 and arm2 lends some support to this (Fig. 6). However, gene conversion prevents us from properly inferring relationships between these LTRs and structural variation in the palindrome by eliminating informative paralogous variations. Inferences about the order of formation of palindromic arms will require analysis in a comparative phylogenetic framework.

### Type A Response Regulator is a sex determination candidate gene

The Y chromosome palindromes in humans contain eight gene families that are expressed predominantly in the testes and which are essential for spermatogenesis [5,38,39]. These genes undergo extensive gene conversion and have high sequence conservation among the copies [5], and they also show evidence of dosage sensitivity [40, 41]. It is therefore reasonable to suspect that the multi-copy genes in the *S. purpurea* palindrome might play important roles in sex determination and/or sex dimorphism. Support for this hypothesis is provided by the type A cytokinin response regulator homologs that occur in palindrome arms 1,2,3a, and 4a (Table 3). The best ortholog of these genes in *P. trichocarpa* is Potri.019G133600. This gene grouped with the Arabidopsis type A response regulators *ARR16* and *ARR17* in the original phylogenetic analysis of this family in *Populus* [42]. Note that this gene was originally designated *PtRR11*, but it is referred to as *RR9* in subsequent publications [43–45], so we will adopt that nomenclature here to avoid confusion. *PtRR9* is expressed primarily in reproductive tissues in *Populus* [42, 45], and is also associated with sex in several *Populus* species [22,43,44]. Further supporting its possible role in sex determination, it was the only gene in the *P. balsamifera* genome that showed clear sex-specific differences in promoter and gene body methylation [43]. This raises the intriguing possibility that this gene provides a shared mechanism of sex determination that plays an essential role in the Chr19 SDR in *Populus* as well as the Chr15 SDR in *Salix*. This is contrary to previous reports that detected no shared sequence composition of these two SDRs, leading to a hypothesis of independent evolution of the sex chromosomes in these genera [20]

The cytokinin signaling pathway has emerged in recent years as a prominent candidate for regulating floral development and sex expression in plants [46, 47]. Cytokinin signaling in Arabidopsis functions as a His-Asp phosphorelay system that transduces the signal from dual function membrane-bound histidine kinases in the endoplasmic reticulum, through histidine phosphotransferases to the type B response regulators (ARR). The type B ARR genes contain both the Asp receiver domain that receives the phosphate from the AHP intermediates, as well as a myb-like DNA binding domain that recognizes conserved motifs in the cytokinin response pathways. The type A response regulators are activated by the Type B ARRs. Type A ARRs contain the receiver domain but lack the DNA binding domain, and are generally negative regulators of the cytokinin response (reviewed in [48]).

In Arabidopsis, disruption of this pathway at multiple stages has resulted in modified reproductive development, with cytokinin typically being implicated in enhancing development of female floral parts [46]. For example, it has recently been shown that the Type B response regulators ARR1 and ARR10 bind to the promoter region of the floral homeotic gene *AGAMOUS*, and down-regulation of these genes inhibits carpel regeneration in an in vitro assay [49]. Similarly, triple mutants of *ARR1*, *ARR10*, and *ARR12* have severely compromised gynoecial development [50]. Proper patterning of the gynoecium is hypothesized to depend upon coordinated expression of genes that are positively regulated by cytokinin in the medial domain of the developing gynoecium, and suppression of these genes in the lateral domains of the gynoecium, potentially by *ARR16* [50]. Intriguingly, *ARR16* is one of the closest orthologs of *PtRR9* [42].

The potential role of cytokinin signaling in dioecy has recently been highlighted by the groundbreaking study by Akagi et al in kiwifruit (*Actinidia spp*) [47]. The authors identified a Type C response regulator (*Shy Girl, SyGI*) on the Y chromosome that was associated with maleness. Overexpression of this gene in Arabidopsis and *Nicotiana tabacum* caused suppression of carpel development, supporting its potential role as a suppressor of female function [14]. This work has some interesting parallels with the results reported here for *Salix* and *Populus*. First, type C response regulators are essentially similar in structure to Type A response regulators, with the main difference being that Type C is not induced by cytokinin. Interestingly, *PtRR9* also was not induced by exogenous cytokinin application [42], though this has not yet been tested with floral tissue. Second, *SyGI* was duplicated from an autosomal gene and subsequently gained a new function on the Y chromosome, much like *SpRR9* has been duplicated from Chr19 in *S. purpurea* and established a distinct pattern of expression, and presumably new functions. However, *RR9* and *SyGI* are clearly not orthologous and likely perform different roles in cytokinin signal transduction. This supports the view that there are numerous ways to achieve separate sexes in plants, and it is likely that a myriad of mechanisms underlie the hundreds of independent occurrences of dioecy in the angiosperms [51], even if a relatively small number of pathways are involved [11, 14].

### Conclusion

We have shown that the SDR of *S. purpurea* contains large palindromic repeats that are similar to the prominent palindromes on mammalian Y chromosomes. We further demonstrated that the *Salix* palindrome is undergoing gene conversion, suggesting some functional similarities to the mammalian sex chromosomes, a striking example of convergent evolution in chromosome structure. We have also demonstrated that the coding sequence undergoing gene conversion in the palindrome, *SpRR9*, is orthologous to a gene that is also associated with sex in *Populus*. This gene is an excellent candidate for controlling sex determination through modulation of the cytokinin signaling pathway. However, much remains to be determined about the underlying mechanism of sex determination. Most importantly, it is currently unclear how the same gene is functioning in an XY system in *Populus* and a ZW system in *Salix*. It is possible that the W chromosome version acts as a dominant promoter of female function, while the Y version is a dominant suppressor of female function, based on the putative roles of cytokinin and the type A response regulators in female development in Arabidopsis. A detailed model should emerge through comparative analysis of the W and Y chromosomes of multiple species in the Salicaceae, which is currently underway.

## Methods

### Initial assembly of the genome

Whole genome assemblies were produced for two *S. purpurea* clones: female clone 94006, and a male offspring of this clone, “Fish Creek” (clone 9882-34), which was derived from a controlled cross between clone 94006 and male *S. purpurea* clone 94001. Clones 94001 and 94006 were collected from naturalized populations in upstate New York, USA. Sequencing reads were collected using the Illumina and PACBIO platforms at the Department of Energy (DOE) Joint Genome Institute (JGI) in Walnut Creek, California and the HudsonAlpha Institute in Huntsville, Alabama. Illumina reads were sequenced using the Illumina HISeq platform, and the PACBIO reads were sequenced using the RS platform. One 400bp insert 2×250 Illumina fragment library was sequenced for total coverage of 183x in clone 94006 and 153x in Fish Creek. Prior to use, Illumina reads were screened for mitochondria, chloroplast, and ΦX174 contamination. Reads composed of >95% simple sequence were removed. Illumina reads <50bp after trimming for adapter and quality (q<20) were removed. For the PACBIO sequencing, a total of 47 P6C4 chips (10 hour movie time) were sequenced for each genome with a p-read yield of 39 Gb and a total coverage of ∼110x per genome (Additional File 1: Table S11). The assembly was performed using FALCON-UNZIP [52]and the resulting sequence was polished using QUIVER [53]. Finally, homozygous SNPs and INDELs were corrected in the release consensus sequence using ∼80x of the 2×250 Illumina reads by aligning the reads using bwa mem and identifying homozygous SNPs and INDELs with the GATK’s UnifiedGenotyper tool [54](Additional File 1: Table S12).

Chromosome-scale assemblies were created using a genetic map derived from 3,697 GBS markers generated for a family of 497 F_2_ progeny from a cross in which the male reference is the father and the female reference is the grandmother. This map is described more completely in a previous publication [55]. This intercross map was used to identify misjoins, characterized by an abrupt change in the *S. purpurea* linkage group. Scaffolds were then oriented, ordered, joined, and numbered using the intercross map and the existing 94006 v1 release assembly [18]. Adjacent alternative haplotypes were identified on the joined contigs, and these regions were then collapsed using the longest common substring between the two haplotypes. Significant telomeric sequence was identified using the (TTTAGGG)_n_ repeat, and care was taken to make sure that it was properly oriented in the production assembly. The remaining scaffolds were screened against bacterial proteins, organelle sequences, GenBank nr and removed if found to be a contaminant. Completeness of the euchromatic portion of the assembly was assessed by aligning *S. purpurea* var 94006 v1 annotated genes to the assemblies. In both cases, 99.7% of the genes were found.

### Identification of W contigs

Contigs derived from the W chromosome are expected to contain some large indels compared to contigs from the Z chromosome due to the lack of recombination between W and Z. These hemizygous regions should exclusively occur in the female specific SDR. To identify these regions, we aligned 2×250 bp Illumina resequencing reads from female clone 94006 and male clone Fish Creek to the new reference using Bowtie2 [56]. Depth of coverage was extracted using samtools-1.2[57]. Median depth was calculated using a non-overlapping sliding window of 10 kb.

To verify if these hemizygous regions are strictly inherited in only female individuals, we used the GBS data from the F_2_ family. GBS reads of 195 offspring of each sex were aligned to the v5 reference with Bowtie2. Due to low coverage and depth of the GBS markers per locus per individual, bam files were merged according to sex in samtools-1.2. Depth was then called in Samtools-1.2 with and max depth was limited to 80,000. Regions continuously covered by GBS reads were defined as GBS intervals. Then, the median of each sex was calculated across all of the intervals. We defined markers as female-specific by integrating the depth from both the F_2_ GBS and 2×250 datasets (restricted to the GBS intervals) using two rules: 1) 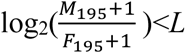, where *L* is the lower bounds of the distribution, defined by the fifth percentile divided by the number of intervals tested (Additional File 2: Figure S6); and 2) 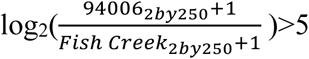.The cutoff for the second criterion was based on the occurrence of a distinct peak in the distribution of the ratios (Additional File 2: Figure S7). Scaffolds that contained at least three female-specific markers were selected as candidate W-female-specific scaffolds (unmapped W pieces). Based on these criteria, only two contigs from the original Chr15 assembly were from W contigs, and the rest were from Z (Additional File 1: Table S5; Additional File 2: Figure S1a).

### Assembly of the Z and W chromosomes

Raw GBS reads used for the original map were demultiplexed and trimmed down to 64 bp for each read by process_radtags (in Stacks 1.44 [58]) with -c -q -r -t 64. Then, trimmed reads of each sequenced individual from the F_2_ family were aligned to the 19 chromosomes and unmapped scaffolds from the main genome and alternative haplotypes from the v4 reference of 94006 using Bowtie 2 [56] with the --very-sensitive flag (-D 20 -R 3 -N 0 -L 20 -i S,1,0.50) to maintain a balance between sensitivity and accuracy. Upon examining the distribution of SNPs in the genome, it became clear that the alternative haplotypes were preventing us from retrieving markers in some regions in the genome, so we repeated the alignments using three different reference sequences: 1) the 19 chromosomes, 2) unmapped scaffolds, and 3) alternative haplotypes. Then, a wrapper script ref_map.pl in Stacks was used to call genotypes with -m 5 (minimum number of reads to create a tag for parents) and -P 3 (minimum number of reads to create a tag for an offspring) on all progeny. Cross type “CP” was chosen since it was the one closest to our cross. Offspring with poor coverage were removed from the downstream analysis.

Once all genotypes were retrieved through Stacks, markers from different loci showing the exact same genotype/segregation across the progeny were binned and only markers from the main genome were kept for mapping. Markers with severe segregation distortion or excessive missing data were excluded, along with twelve offspring with very low call rate. Genotypes were imputed and corrected based on inferring haplotypes in the two F1 parents from segregation of the markers in the progeny.

The grandparents of the F_2_ cross have extensive stretches of shared haplotypes, possibly due to historic inbreeding in this naturalized population. This results in long runs of heterozygosity and homozygosity in the F_1_ progeny. This inhibits integration of backcross and intercross markers by available mapping algorithms like those in the Onemap package [59]. To circumvent this problem, all intercross markers were translated to female and male backcross markers by identifying the parental origins of alleles based on parental phases and physical position in the assembly. Also, putatively hemizygous markers were recoded as backcross markers using sequence depth to infer genotypes. For example, markers with the segregation pattern +/- x -/- were recoded as AB x BB. These genotypes were also imputed and corrected based on the inferred haplotypes of the two F1 parents.

Onemap v2.1.1 was used to form initial linkage groups. For each chromosome, there are two phased linkage groups from each backcross type. However, this phase information derived from the F_2_ family is only for the F1 parents, which cannot be directly used for phasing haplotypes in the grandmother, clone 94006. By comparing parental genotypes from one LG to those of the grandparents, we inferred which of the 94006 haplotypes were inherited by each F_1_. These results were used as a piece of evidence for identifying W-linked scaffolds/contigs, as well as estimating the overall occurrence of chimeric contigs in the assembly. After building a framework genetic map using markers from the main genome, non-distorted markers from unmapped main scaffolds and alternative scaffolds were added.

All unmapped scaffolds were manually checked to see if they matched the phase information or contained female-specific markers. Those that were identified as Z scaffolds/contigs were excluded from the W map. The new W and Z were assembled using the python package ALLMAPS [60] to order and orient scaffolds and reconstruct chromosomes based on the genetic map. Only the order of the female backcross map was used to assemble the W, and ALLMAPS was set not to break contigs. This new map-based assembly containing two versions of chromosome 15 (Chr15Z and Chr15W) is version 5 of the *S. purpurea* var. 94006 genome.

To identify Z-degenerate regions and insertions in the female specific W (FSW), we realigned the 2×250 reads of 94006 and Fish Creek to the 94006 v5 reference as described above, except we removed Chr15Z out of the reference. Depth was calculated using samtools, and the median depth of 50kb non-overlapping windows across was calculated with an in-house script. Regions where medians of Fish Creek depth are no greater than 10 were considered as insertions in the FSW, and regions with greater depth were considered Z-degenerate regions. This analysis was repeated with a 10 kb window as well to enhance the resolution of the analysis.

### Annotation of the genome

Transcript assemblies were constructed from ∼126M pairs of 2×76bp (94006) or 2×150bp (Fish Creek) paired-end Illumina RNA-seq reads using PERTRAN. 188,628 transcript assemblies were constructed using PASA from the RNA-seq transcript assemblies. Loci were determined by transcript assembly alignments and/or EXONERATE alignments of proteins from *Arabidopsis thaliana*, soybean, poplar, cassava, brachypodium, grape, and Swiss-Prot proteomes, and high confidence *Salix purpurea* Fish Creek gene model peptides to repeat-soft-masked *S. purpurea* 94006 genome using RepeatMasker with up to 2K BP extension on both ends unless extending into another locus on the same strand. Gene models were predicted by homology-based predictors, FGENESH+, FGENESH_EST (similar to FGENESH+, EST as splice site and intron input instead of protein/translated ORF), and EXONERATE, PASA assembly ORFs (in-house homology constrained ORF finder) and from AUGUSTUS via BRAKER1. The best scored predictions for each locus were selected using multiple positive factors including EST and protein support, and one negative factor: overlap with repeats. The selected gene predictions were improved by PASA. Improvement includes adding UTRs, splicing correction, and adding alternative transcripts. PASA-improved gene model proteins were subject to protein homology analysis to the above mentioned proteomes to obtain Cscore and protein coverage. Cscore is a protein BLASTP score ratio to MBH (mutual best hit) BLASTP score and protein coverage is the highest percentage of protein aligned to the best homolog. PASA-improved transcripts were selected based on Cscore, protein coverage, EST coverage, and its CDS overlapping with repeats. The transcripts were selected if its Cscore is larger than or equal to 0.5 and protein coverage larger than or equal to 0.5, or it has EST coverage, but its CDS overlapping with repeats is less than 20%. For gene models whose CDS overlaps with repeats for more than 20%, its Cscore must be at least 0.9 and homology coverage at least 70% to be selected. The selected gene models were subject to Pfam analysis and gene models whose protein is more than 30% in Pfam TE domains were removed. Incomplete gene models, low homology supported without fully transcriptome supported gene models and short single exon (< 300 BP CDS) without protein domain, or good expression gene models were manually filtered out.

To annotate potential genes or coding regions in the palindrome that were missed by the automated annotation, the full nucleotide sequence of arm1 (about 20 kb) was submitted to the Fgenesh online service (http://www.softberry.com/berry.phtml?topic=fgenesh) with specific gene-finding parameters for *Populus trichocarpa*. The predicted peptide sequences were searched against predicted proteins from *Populus trichocarpa* v3.0, and *Arabidopsis thaliana* TAIR10 in Phytozome 12 (https://phytozome.jgi.doe.gov/) to find the closest homologous annotation. The protein domains were identified using hmmscan in HMMER (v3.1b1, http://hmmer.org/) against the Pfam-A domains (release 32, https://pfam.xfam.org).

### Comparison of Z and W orthologous genes

Allelic genes on the Z and W chromosomes were identified by performing a reciprocal blastp of all primary annotated peptide sequences in the main genome with default parameters. Mutual best hits were identified with over 95% identity over at least 80% of the transcript. Tandem duplications were identified as genes with expectation values of 1×10^-10^ that occurred within a 500 kb window. In these cases, one representative gene from each tandem array was used as a representative sequence, and the mutual best hit outside the tandem array was identified as above.

To identify allelic gene pairs for calculation of synonymous substitutions between the Z and W alleles, a reciprocal blast of all primary annotated peptide sequences was run with “blastall –p blastp -i -e 1e-20 -b 5 -v 5 -m 8”, and MCscanX was run with default parameters [61]. The synonymous and nonsynonymous substitution rate of each gene pair in each syntenic block was estimated by aligning the sequences with CLUSTALW [62] and using the calc_all_KaKs_pairs script in MCscanX. Only pairs between Chr15W and Chr15Z, and scaffold_844 were used for estimating the divergence between Z and W.

### Identification of sex-associated loci

Loci associated with sex were identified using 60 non-clonal individuals from a naturalized population of *S. purpurea* [63]. GBS reads from each individual were aligned to the 94006v5 genome without Chr15Z using Bowtie2. Genotypes were called in Stacks 1.14 using the ref_map.pl wrapper and the populations module with a minimum minor allele frequency of 0.1 and a genotyping rate of 0.1. Loci with greater than 40% missing data were removed. Association with sex was performed using emmax [64] as described previously [18].

### Detection of palindromic repeats

We conducted a systematic scan of large highly identical inverted regions. Each chromosome and scaffold in the main genome was aligned to itself with LASTZ 1.03.66 with the following flags: --gapped --exact=100 --step=20. An exact value of 100 means that 100-base exact match is required to qualify as an High Scoring Pair in the output. Using a 1-kb sliding window and 100-bp steps, an intrachromosomal hit was kept if it matched the following criteria: 1) the hit has the highest identity in the window, 2) greater than 2 kb in length, 3) aligned regions are not further apart than 500 kb, and 4) there is a corresponding hit on the negative strand. Candidate palindromes also satisfied the following criteria: 1) identity greater than 99.0%, 2) the total number of hits from both strands should be less than 4, with at least one on the negative strand, and 3) large inverted hits are at least 8 kb in length. Each region was manually checked and assembly artifacts (e.g., corresponding to contig boundaries) and complex repetitive regions were excluded.

Paralogous gene copies on autosomes were retrieved from the reciprocal blastp results described above. Paralogous genes within the palindrome arms were aligned along with paralogous copies from the autosomes using Muscle using default parameters provided in MEGA 5. In a few cases, the resulting alignments were adjusted manually (Supplemental Materials: AdditionalFile3). A Neighbor-Joining tree with default parameters was built using MEGA 5 [65].

To identify recent insertions of transposable elements within the palindrome, LTRharvest [66] was run with the sequence of the palindromic on Chr15W from 8,778 kb to 9,015 kb with the target site duplication restricted to 5 bp to 20 bp. To find the protein domains in the coding region, a protein domain search against Pfam-A domains (release 32) was performed using the hidden Markov model methods implemented in LTRdigest (–hmms flag) [67]. Predicted LTR retrotransposons were determined to be non-automonous when coding regions did not contain any *gag* or *pol* related domains.

To estimate the substitution rate between the flanking LTR repeats, 5’ and 3’ repeats of each LTR retrotransposon predicted from LTRharvest were aligned by MUSCLE using default parameters provided in MEGA 5. After all gaps were removed, both number of differences and substitution rate were estimated in MEGA5. For number of differences, transitions and transversions were both included with a uniform rate. Substitution rate was modeled using the Kimura 2-parameter model provided in MEGA5, and the rate variation among sites was modeled with a gamma distribution (shape parameter = 1).

### Detection of gene conversion

As evidence of gene conversion, we searched for regions that were differentiated between species but concordant among the palindrome arms. To accomplish this, we aligned paired-end reads from a female clone of *S. suchowensis* (srx1561933) to the 94006 v5 female reference, plus alternative haplotypes. This yielded an 82.9% overall alignment rate on average. Mis-mapped reads originating from the autosomes were manually identified by scrutinizing the alignments, and only reads that mapped exclusively to the palindromic regions were retained. These reads were re-aligned to Arm1. SNPs and indels were called using mpileup with a minimum mapping quality of 20 and depth less than 300.

### Expression Profiling

RNAseq data was obtained from catkins of 10 female and 10 male F_2_ progeny. RNAseq data was also obtained from multiple tissues of clones 94006 and Fish Creek. All sequences were Illumina 2×150 bp reads, except for 94006, which were 2×76 bp reads. Transcripts from the palindrome can be highly identical among arms and with other paralogous sequences on the autosomes, which can complicate estimation of gene expression. Thus, all predicted coding sequences from the same gene family in the palindrome were aligned to the autosomal paralogs, and conserved sequences were masked in the reference genome. Salmon-0.11.3 [68] was used to quantify (salmon quant) the raw read count for each sample mentioned above with the gcBias flag as suggested by the developers. Heatmaps were generated separately for each group of palindrome genes, using log_2_ transformed data normalized with respect to library size or by variance stabilizing transformations (VST) using the R packages pheatmap and Deseq2 [69].

## Availability of Data and Materials

All sequence data used in this manuscript have been deposited in the NCBI Sequence Read Archive (https://www.ncbi.nlm.nih.gov/sra). Accession numbers are available in Additional File 1: Table S13. The genome assemblies and annotations are available through Phytozome (https://phytozome-next.jgi.doe.gov).

## Acknowledgements

We thank Fred Gouker and the teams at the New York State Agricultural Experiment Station and West Virginia University for help with management and phenotyping of the field trials used in this study. We also thank the teams at Phytozome and JGI for facilitating public distribution of the data. We also thank Niels Müller and Mathias Fladung for helpful discussions.

## Author Contributions

R.Z. and D.M.-S. assembled and analyzed the W chromosome; J.S. and J.W.J. assembled the genome; D.K., A.S., L.S., G.A.T., C.C., and K.B. sequenced the genome and transcriptomes; S.S. annotated the genome; S.D., L.B.S., T.M., J.L., and M.O. designed and led the study. S.D, R.Z., and D.M.-S. wrote the manuscript.

