## Supplemental Figures for "A Willow Sex Chromosome Reveals Convergent Evolution of Complex Palindromic Repeats"

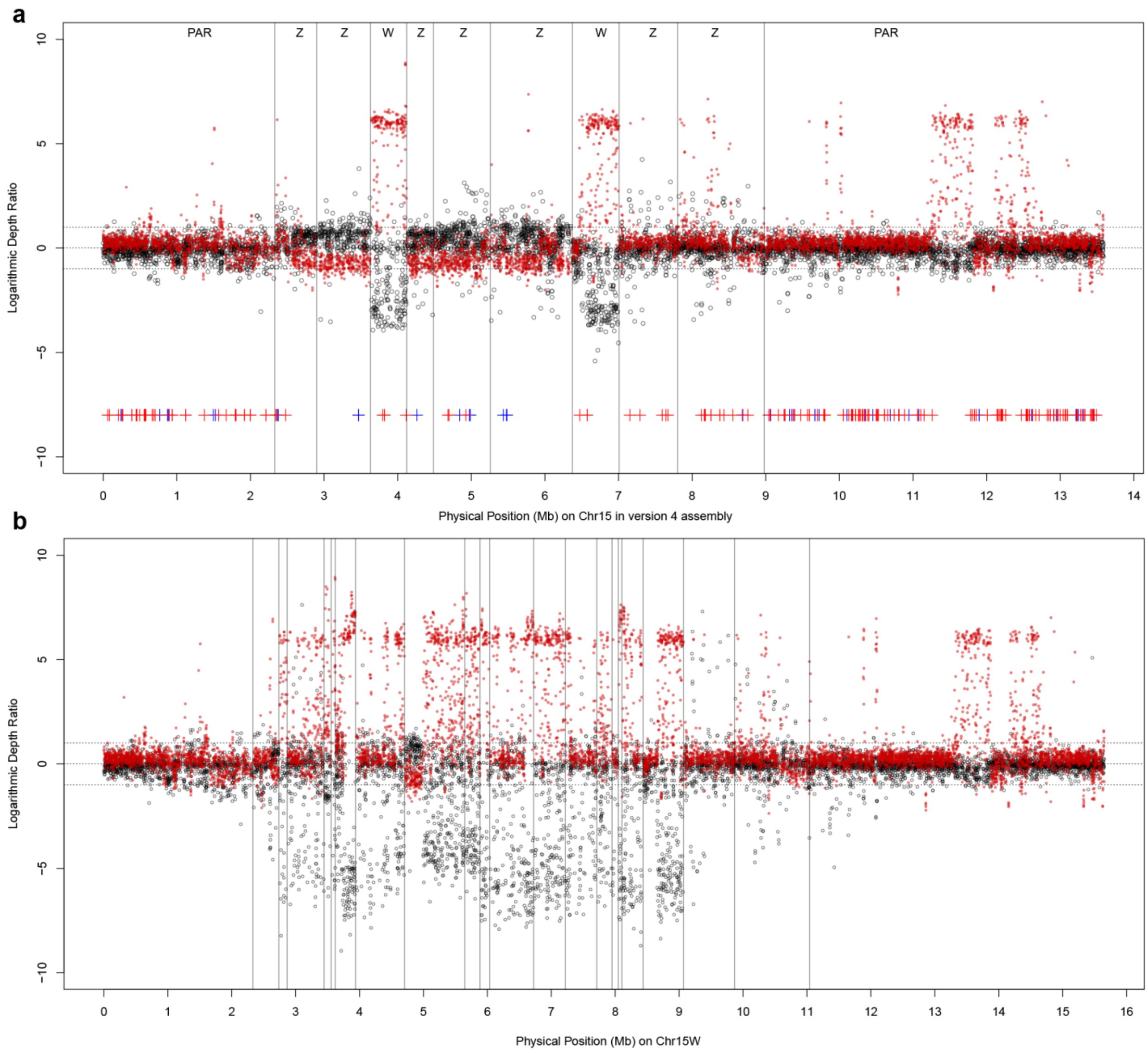


**Figure S1 Sex-specific depth of Chr15 for two assemblies of female clone 94006. a**. Initial assembly guided by mapped SNP markers only. Boundaries of contigs are represented with vertical lines. Each contig is categorized as pseudo-autosomal region (PAR), Z, or W according to the logarithmic depth ratio between female and male sequence alignments. Ratios of GBS marker depth ($\log_2(M/F)$) for 200 progeny from an F2 pedigree (family 317) are shown by black dots, and ratios of the two reference individuals from Illumina 2x250 resequencing reads $\log_2(F/M)$ are shown by red dots. Near the bottom, red-crosses represent markers that are inherited from the female parent, and blue crosses are markers inherited from the male parent. **b**. Chromosome 15W, following reassembly using scaffolds with female-specific alleles and/or female-biased depth ratios.

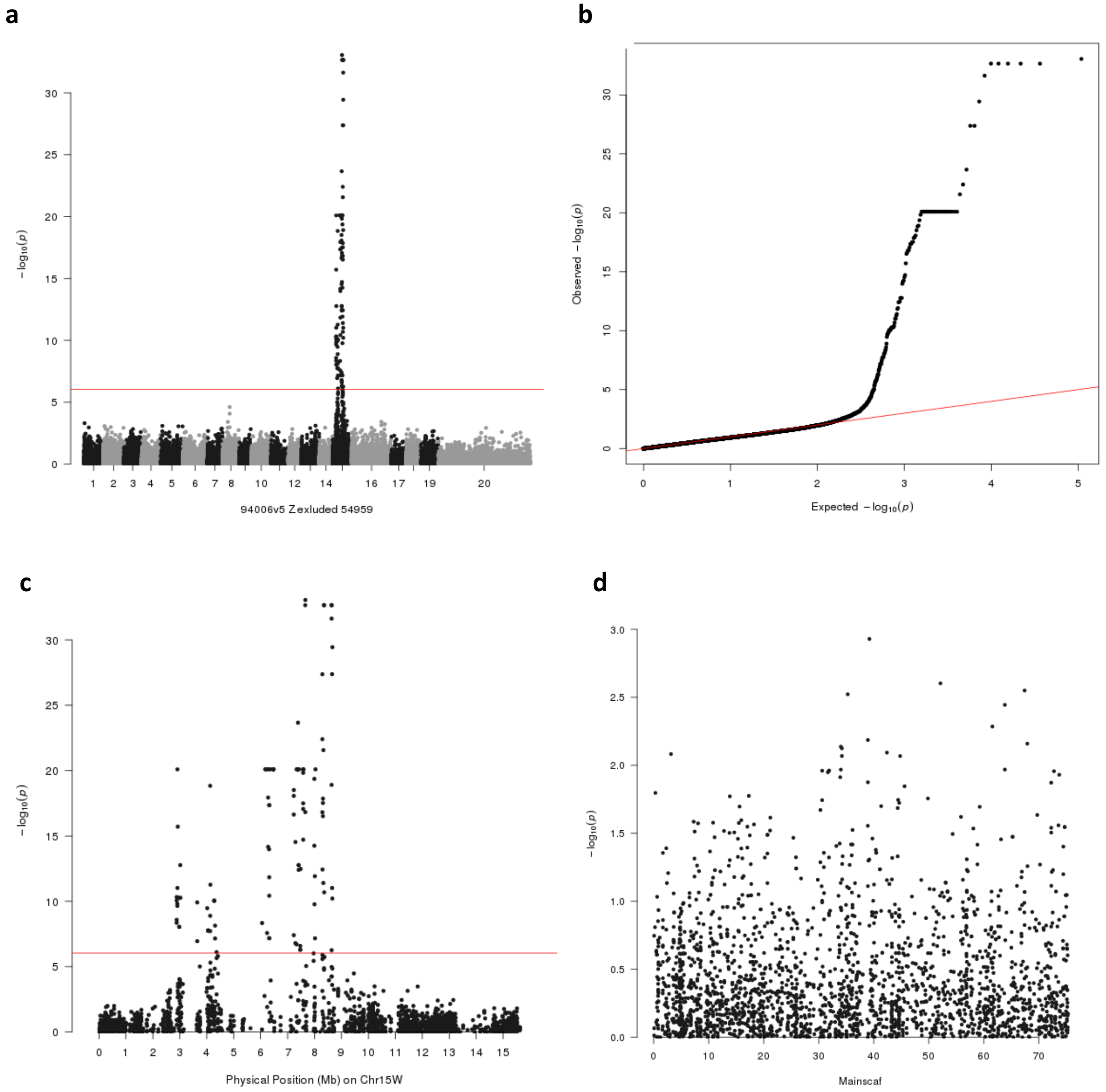


**Figure S2 Association of sex with new 94006v5 assembly. a**. Manhattan plots showing association of sex repeated with the v5 genome assembly, with Chr15Z removed. The analysis was performed with a natural population of 60 non-clonal individuals. The red line indicates a Bonferroni cutoff 9.10 x $10^{-7}$ with 54,959 tested SNPs. **b**. QQ-plot for the association analysis. **c**. Manhattan plot for Chromosome 15W. **d**. Manhattan plot for unplaced scaffolds from the main genome. None of these showed significant association with sex.

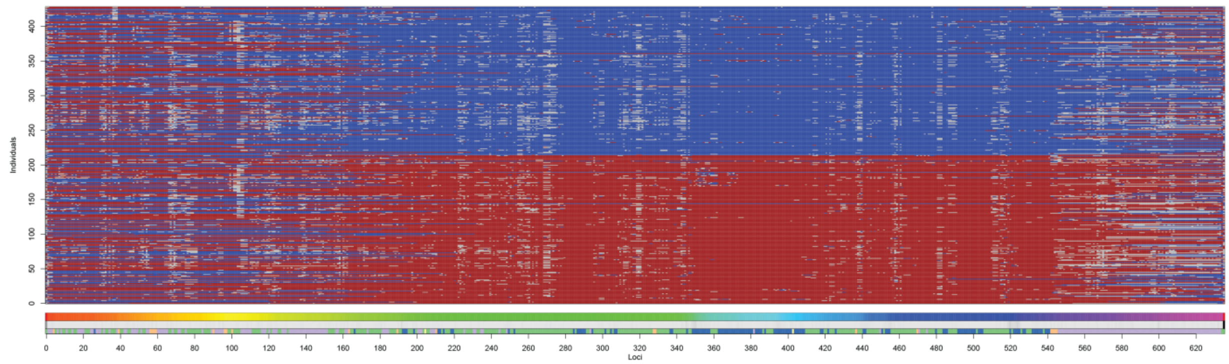

**Figure S3 Recombination among parental haplotypes in F2 progeny.** GBS markers are ordered along the genetic map. 214 $F_2$ progeny from each sex are in the rows, with males at the top of the figure. Red cells represent alleles derived from the W haplotype, and blue cells represent the Z haplotype according to the maternal (Wolcott) genetic map. Gray cells represent missing data that could not be imputed.

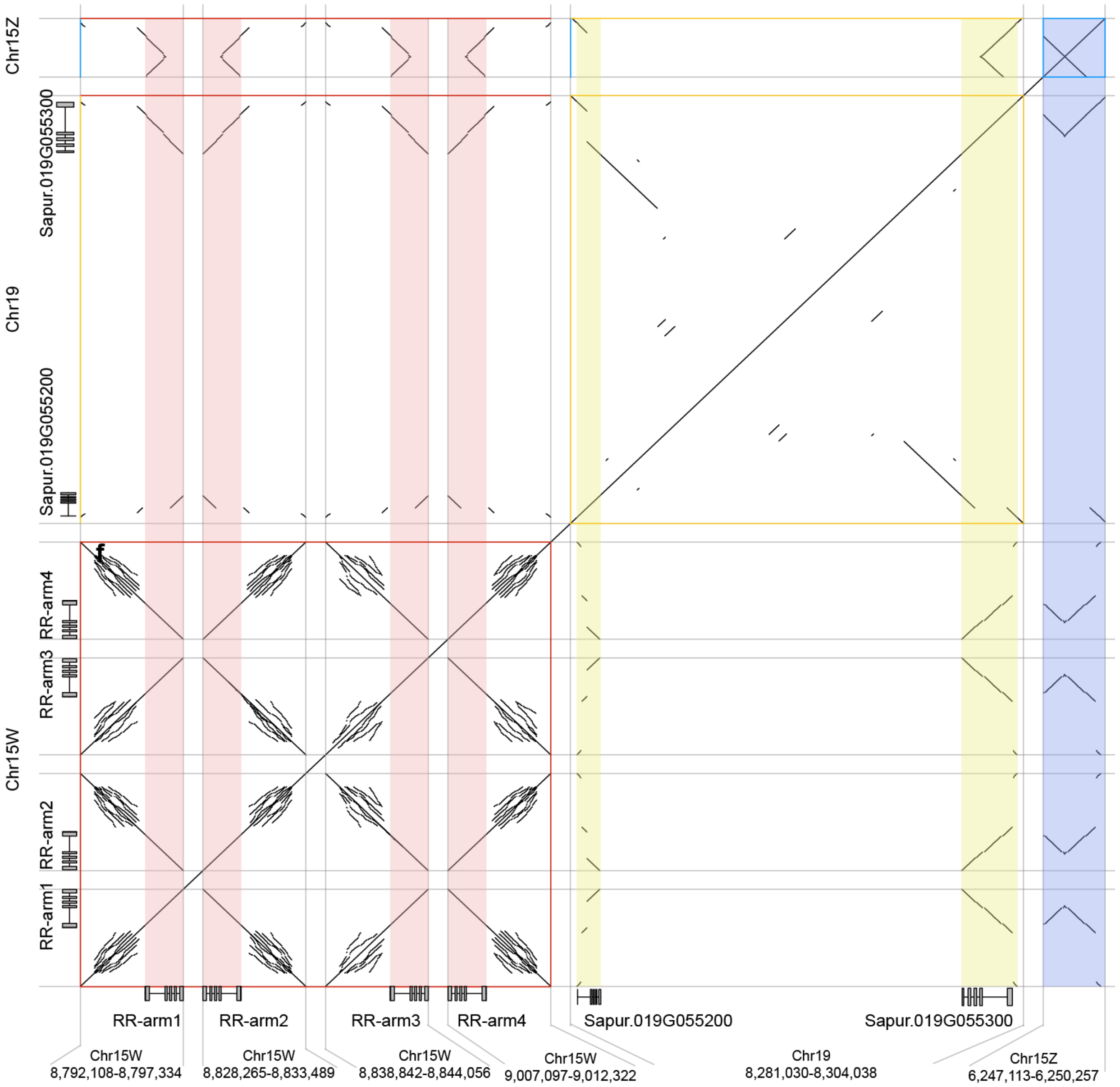


**Figure S4 Dotplots of regions containing portions of the RR gene.** This dot plot was generated from data produced by aligning sequences from identified regions containing the RR genes (complete or partial) on Chr15W palindrome (red), Chr15Z (blue), and Chr19 (yellow) using LASTZ. Colored shading indicates the X axis location of genes and genes models, which are also displayed on both axes. Notice that the Chr15Z block (blue) contains a truncated portion of the gene, and was not annotated. Three colored squares along the diagonal line show the palindromic structures. Horizontal and vertical lines with different colors indicate the area of pairwise alignments between RR genes from different chromosomes.

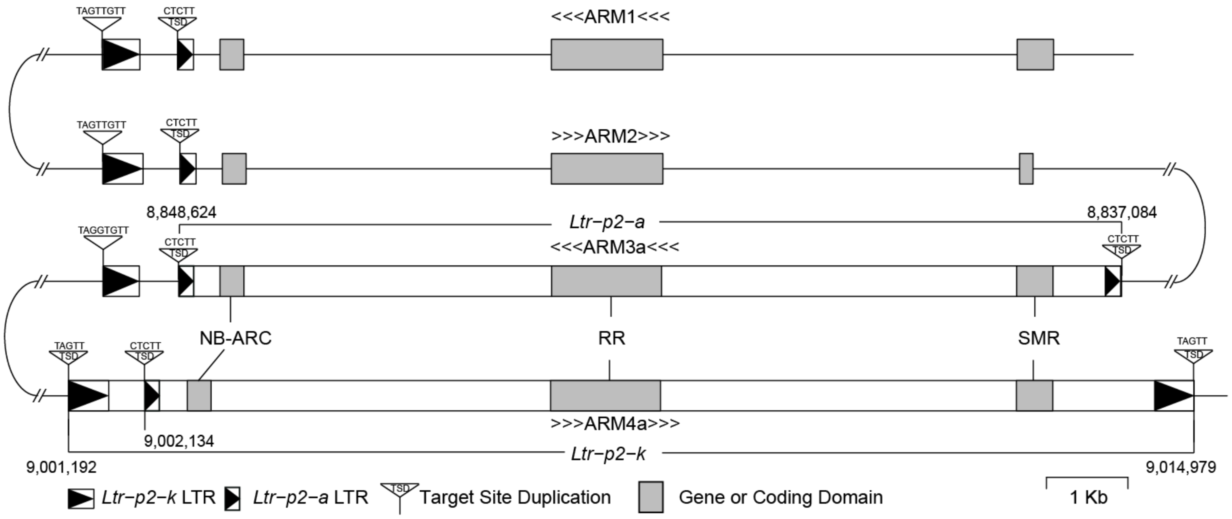


**Figure S5 Arrangement of *Ltr-p2-a* and *Ltr-p2-k* in the palindrome.** Two non-autonomous LTR retrotransposons on arm3 and arm4 are shown with their target site duplication (TSD) sequences, long terminal repeats (LTRs), and genes or domains highlighted. Duplicated sequence features are also labeled on arm1 and arm2. Numbers indicate the coordinates of these transposable elements on the W chromosome.

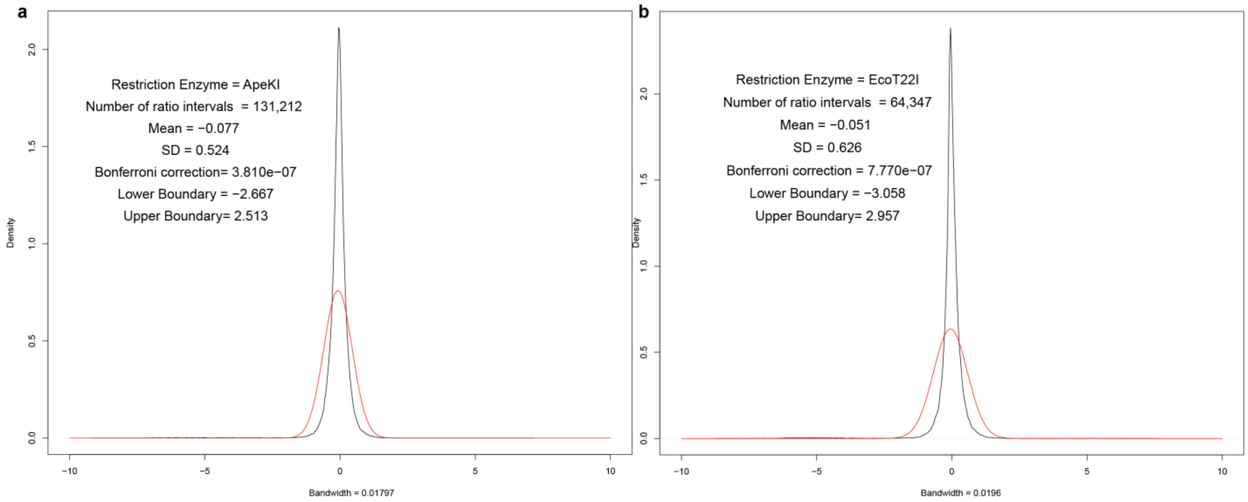


**Figure S6 Distribution of depth ratios (**$\log_2(\frac{M_{195}+1}{F_{195}+1})$**) of GBS reads aligned to the female 94006 v4 genome**. The distribution of depth ratios of GBS markers of *Ape*KI is indicated by the black line, and normal distribution with the same mean and standard deviation (SD) is indicated with a red line. To detect outliers, such as intervals only covered in one sex, lower and upper boundaries were determined according to the Bonferroni corrected percentile (0.05/number of intervals) of this normal distribution. **b**. Same process was applied with the GBS markers that were generated from *Eco*T22I.

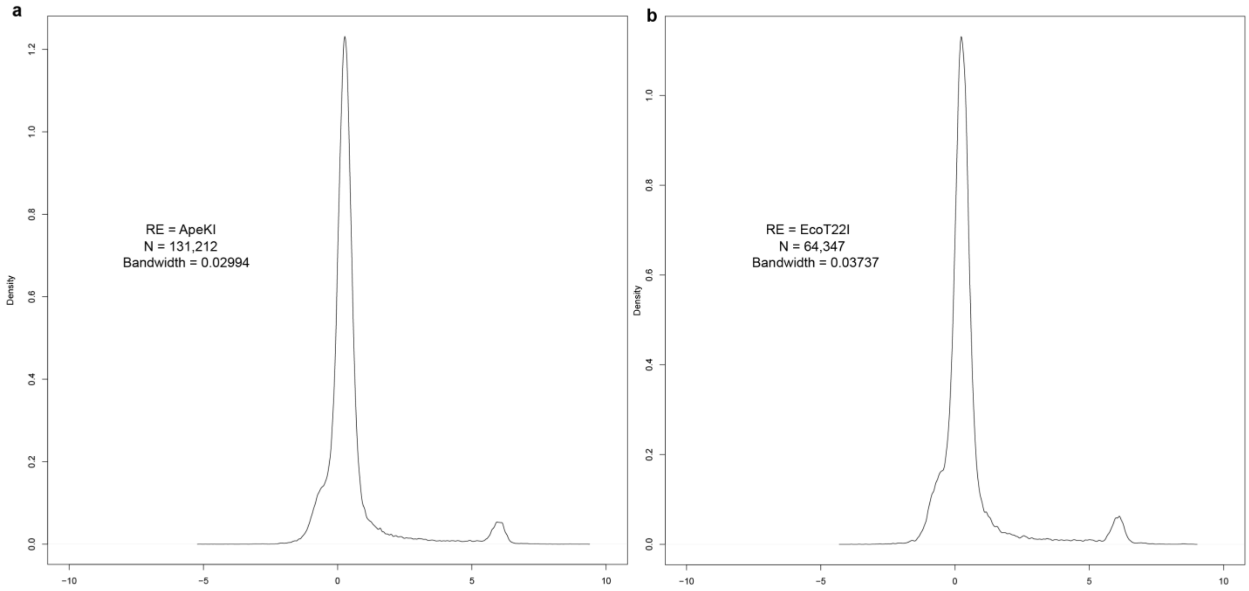


**Figure S7 Distribution of depth ratios ($\log_2(\frac{94006_{2by250}+1}{Fish\ Creek_{2by250}+1})$) of Illumina reads aligned to the female 94006 v4 genome. a.** Counts are only from intervals defined by GBS markers from the $F_2$ family to facilitate comparisons. The peak around 6 putatively represents sequences derived from the W chromosome, as well as deletions in Fish Creek relative to 94006. **b**. Same process was applied with the GBS markers that were generated from *Eco*T22I.
